# Neural signatures of pathological hyperdirect pathway activity in Parkinson’s disease

**DOI:** 10.1101/2020.06.11.146886

**Authors:** Ashwini Oswal, Chien-Hung Yeh, Wolf-Julian Neumann, James Gratwicke, Harith Akram, Andreas Horn, Ludvic Zrinzo, Tom Foltynie, Patricia Limousin, Rafal Bogacz, Masud Husain, Peter Brown, Vladimir Litvak

## Abstract

Parkinson’s disease (PD) is characterised by the emergence of pathological patterns of oscillatory synchronisation across the cortico-basal-ganglia circuit. The relationship between anatomical connectivity and oscillatory synchronisation within this system remains poorly understood. We address this by integrating evidence from invasive electrophysiology, magnetoencephalography, tractography and computational modelling in patients. Coupling between supplementary motor area (SMA) and subthalamic nucleus (STN) within the high beta frequency (21-30 Hz) range correlated with fibre tract densities between these two structures. Additionally within the STN, non-linear waveform features suggestive of cortical synchronisation correlated with cortico-STN fibre densities. Finally, computational modelling revealed that exaggerated hyperdirect cortical inputs to the STN in the upper beta frequency range can provoke the generation of widespread pathological synchrony at lower beta (13-20 Hz) frequencies. These observations reveal a spectral signature of the hyperdirect pathway at high beta frequencies and provide evidence for its pathophysiological role in oscillatory network dysfunction in PD.

**One sentence summary:** Signatures of the hyperdirect pathway and its likely role in pathological network disruption in Parkinson’s disease are identified.

## Introduction

Parkinson’s disease is a common disorder of movement, which is characterised by nigrostriatal dopamine depletion and the emergence of stereotyped patterns of oscillatory synchronisation within cortico-basal-ganglia circuits. Excessive synchronisation across beta band frequencies (13-30 Hz) is a hallmark of the Parkinsonian dopamine depleted state. By simultaneously recording cortical activity with EEG or magnetoencephalography (MEG) and intracranial local field potentials (LFP) in patients undergoing surgery for the insertion of Deep Brain Stimulation (DBS) electrodes it is possible to explore patterns of long range synchronisation that emerge within cortico-basal-ganglia circuits ^1–3^. Using this approach, it has been previously shown that the STN couples with motor/premotor activity at beta frequencies, with the cortex predominantly driving STN activity ^2,4^.

In contemporary models of the basal-ganglia-thalamocortical loop, cortical activity is thought to be transmitted to subcortical regions by three streams – the hyperdirect, direct and indirect pathways – which act in conjunction to shape the dynamics of action initiation and selection. The direct and indirect pathways provide cortical inputs to basal ganglia via the striatum ^5–8^. The hyperdirect pathway is a monosynaptic axonal connection, thought to be at least partly formed from axon collaterals of corticobulbar and corticospinal fibres, which runs from frontal cortex to the STN and is proposed to provide rapid inhibition for action suppression ^9–11^. The physiological properties of the hyperdirect pathway have been studied in humans and animals using a combination of techniques including tracer studies ^12^, evaluation of evoked responses to DBS ^11,13,14^ and non-invasive tractography ^10,15,16^.

An improved understanding of the relationship between cortico-basal-ganglia anatomical projections and the generation of beta band oscillatory synchrony is essential to fulfil a critical gap in our understanding of network dysfunction in PD and could inform the development of more spatially and temporally patterned DBS therapies ^17^. In this regard, previous studies have speculated on the potential importance of an exaggerated hyperdirect pathway ^4,18–21^, but details regarding pathophysiological mechanisms are lacking. Previous work demonstrates that beta band coupling between the cortex and STN is segregated such that mesial motor/premotor areas drive STN activity at high beta frequencies (21-30 Hz) ^2,4^. Synchrony at lower beta frequencies (13-21 Hz) is also detectible in the parkinsonian STN and is considered to be more directly pathological as it is suppressed by both DBS and L-dopa therapy ^22–24^ with some studies also demonstrating a correlation between the extent of treatment related beta suppression and clinical improvement ^4,25–28^. This leads to the hypothesis that cortical coupling with the STN at high beta frequencies may reflect hyperdirect pathway activity ^4^. If high beta frequencies are reflective of the hyperdirect pathway, synchrony within this frequency range should be relatively restricted to the STN and its interactions with cortex. Furthermore, we would predict an overlap between cortico-STN *anatomical* connectivity of the hyperdirect pathway and the profile of cortico-STN *functional* connectivity at high beta frequencies. Moreover, STN beta waveform features suggestive of synchronisation ^29,30^ to a cortical source would also be expected to correlate with the extent of cortico-STN anatomical *connectivity*.

To test these hypotheses, we performed simultaneous MEG and intracranial LFP recordings in Parkinsonian patients undergoing surgery for the insertion of DBS electrodes in either the STN or the internal segment of the globus pallidus (GPi). Synchrony profiles of the STN and GPi networks were compared. Cortico-STN *anatomical* connectivity derived from individual patient electrode localisations and open source tractography connectomes was integrated with individual patient MEG and LFP derived cortico-STN *functional* connectivity in order to establish the relationship between hyperdirect pathway fibre density and cortico-STN coupling at high beta frequencies.

Our findings indicate that cortical connectivity with the STN at high beta frequencies reflects activity within the hyperdirect pathway. Our empirical findings are recapitulated by a biophysical model which additionally reveals that an exaggerated hyperdirect pathway in PD may lead to the generation of subcortical synchrony at lower beta frequencies (13-21 Hz) which are considered to be more directly pathological.

## Results

### Differences in local synchrony between the STN and GPi at high beta frequencies

**Figure 1** shows trajectories and contact locations for all electrodes targeting the STN (blue) and GPi (red) in MNI space. Contact localisation for individual patients is shown in **Supplementary Figures 1 and 2**. **Figure 2A** depicts the mean spectral power for all STN and GPi bipolar contacts, whilst **Figure 2B** reveals the mean spectral power of only the oscillatory component (after removing the 1/f component) for the same STN and GPi contacts. In both structures there is a peak centred at approximately 7-8 Hz. Subjects also displayed oscillatory peaks within the low (13-21 Hz) and high beta (21-30 Hz) frequency ranges. Although oscillatory power at low beta frequencies (13-21 Hz) was not significantly different between the two structures, we did observe oscillatory power differences between 24 and 34 Hz (**Figure 2B**; peak *t* = 5.7, FWE *p* < 10^−4^), encompassing the high beta frequency range, such that power over this range was greater within the STN than within the GPi.

**Fig 1.**
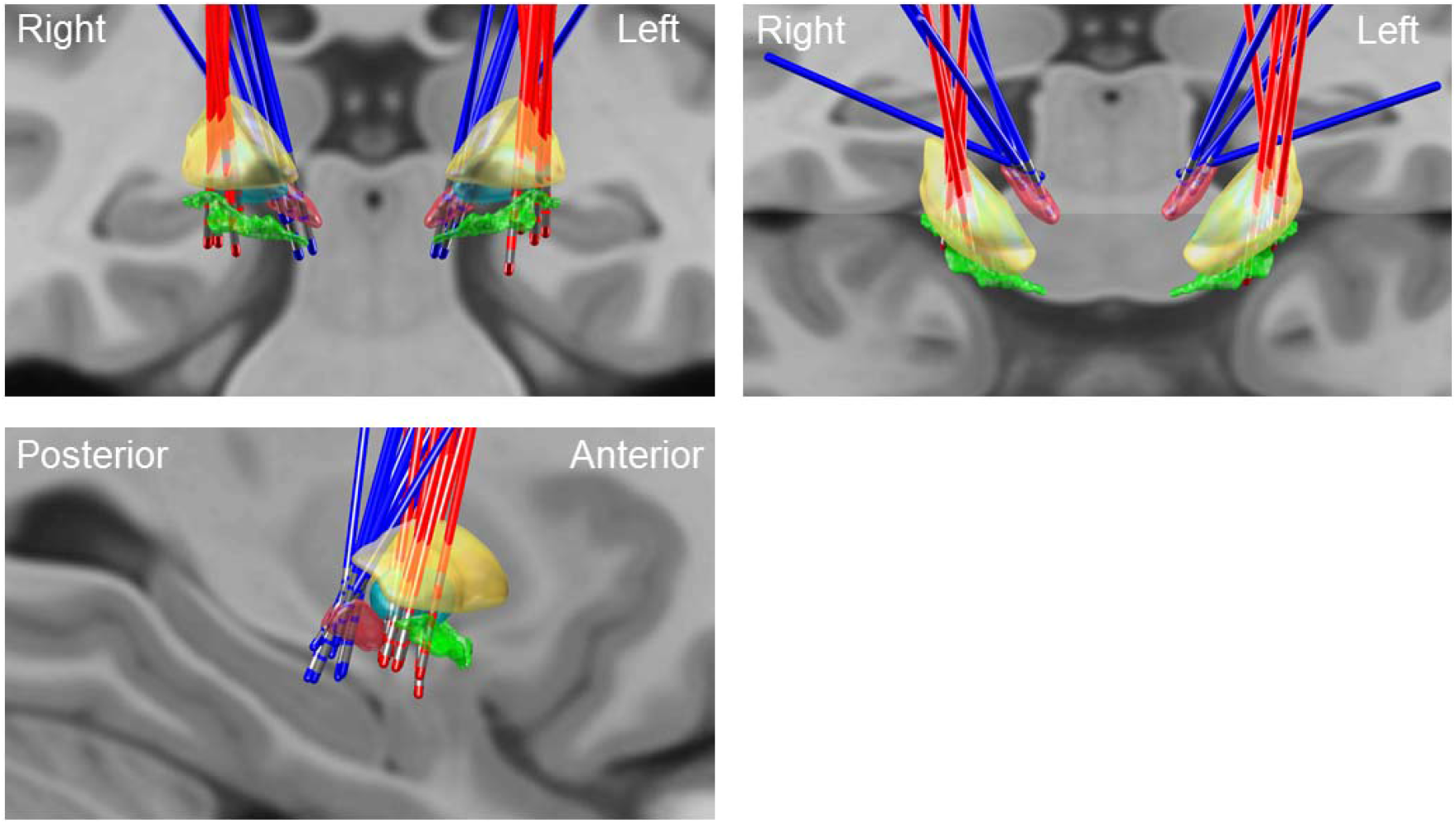
Localisation of electrodes in MNI space. Electrodes targeting the STN are coloured blue whilst electrodes targeting the GPi are coloured red. Templates of the STN (red), GPi (turquoise), GPe (yellow) and nucleus basalis of Meynert (NBM; green) are superimposed on a T1-weighted structural MRI for visualisation. The 3D image is viewed in the sagittal, coronal and angled (45 degree) axial planes.

**Fig 2.**
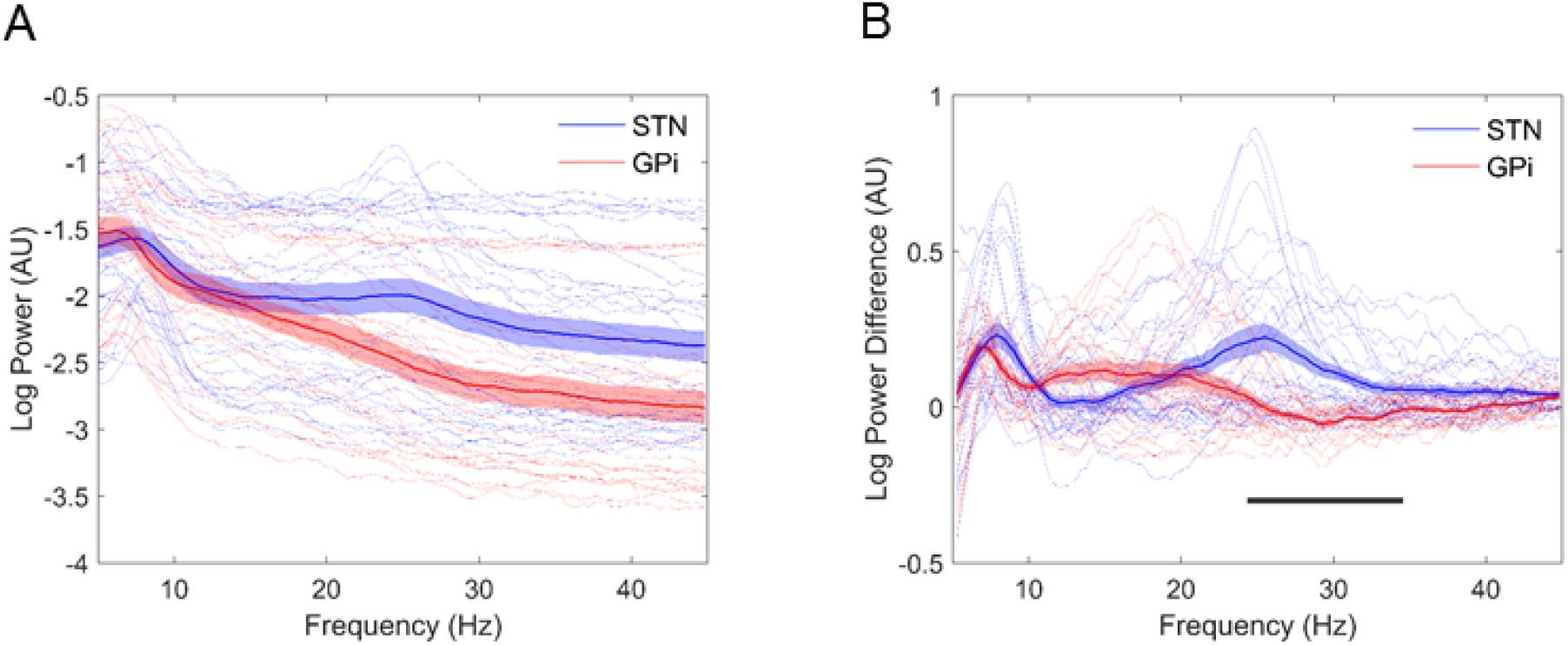
Local synchrony distinguishes the STN and GPi. Mean log spectral power for all STN (thick blue line) and GPi (thick red line) contacts is plotted in (A). Panel (B) displays the mean log power of the oscillatory component after subtraction of the non-oscillatory 1/f component. Spectral profiles of individual STN and GPi contacts are also shown in both panels by the feint blue and red lines respectively. In both structures there is a peak centred at approximately 7-8 Hz. Subjects also displayed oscillatory peaks within the low (13-21 Hz) and high beta (21-30 Hz) frequency ranges. Oscillatory power between 24 Hz and 34 Hz (indicated by the black line in (B)), encompassing the high beta frequency range, was significantly greater in the STN than in the GPi.

### Cortical-subcortical coherence at high beta frequencies occurs preferentially within the cortico-STN network

Comparison of the profile of cortical coherence for the STN and GPi, revealed an interaction between frequency band and electrode location such that specific cortical regions displayed greater coherence with the STN than they did with the GPi, preferentially at high rather than at low beta frequencies. Regions displaying this interaction included the SMA and mesial areas of primary motor cortex (**Figure 3A** upper panel; peak *t* = 4.5, FWE p < 3×10^−3^ at MNI co-ordinates 8 -46 82). In **Figure 3A** the blue and green contour lines represent the boundaries of primary motor cortex and the SMA derived from the Automated Anatomical Labelling (AAL) atlas. Interestingly the interaction effect did not extend as far laterally as the hand area of primary motor cortex (see lower panel of **Figure 3A**). In contrast there was no main effect of frequency band or electrode location (peak *t* = 1.8 and 1.9 respectively, in both cases FWE p > 0.1).

**Fig 3.**
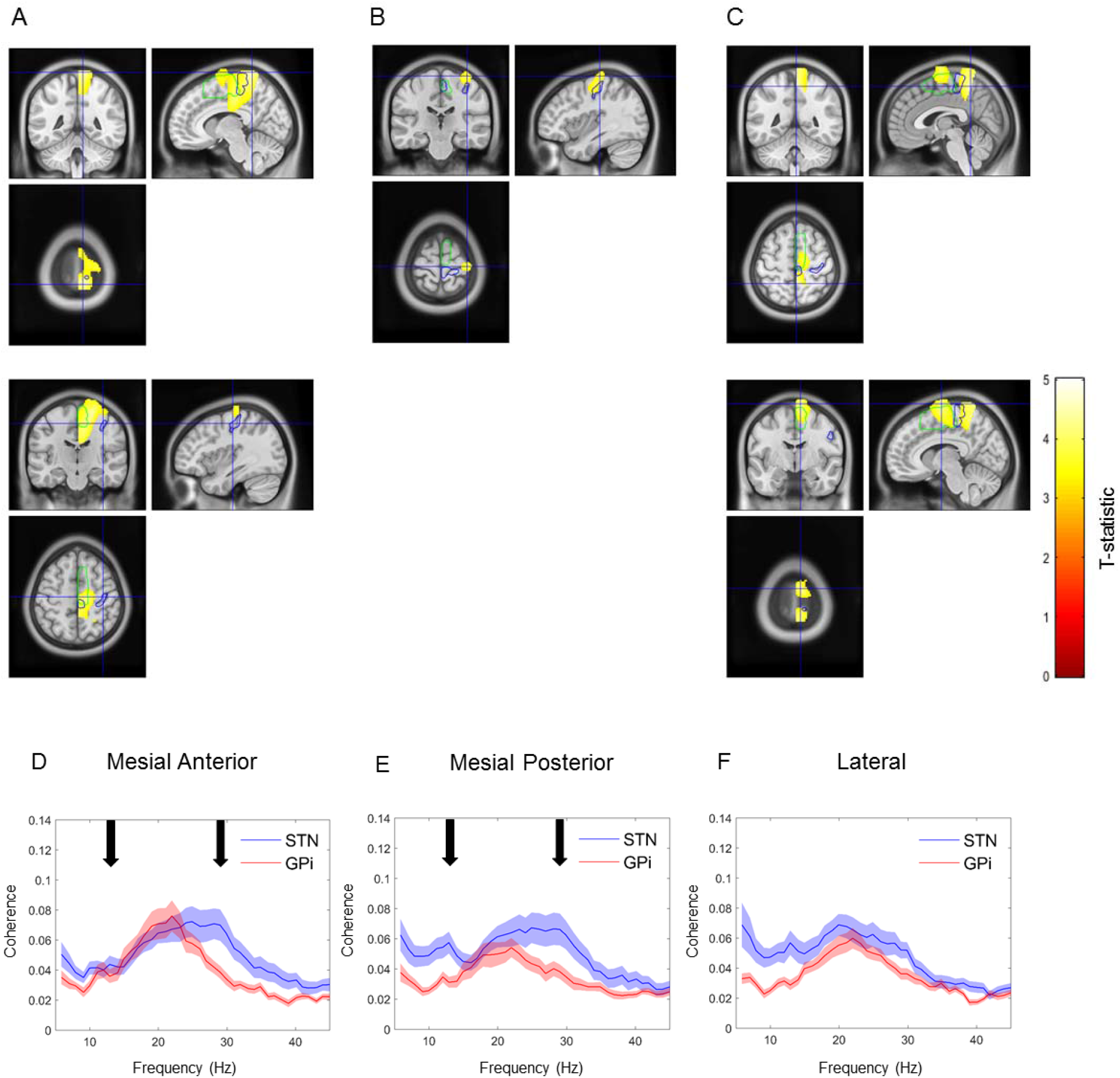
Differences in cortico-STN and cortico-GPi coherence; high beta band coherence segregates to the cortico-STN network. SPMs showing results of the 2x2 ANOVA with factors frequency (low beta vs high beta) and target nucleus (STN vs GPi). T-values of voxels within significant clusters are superimposed onto a T1-weighted MRI with the colour bar indicating the value of the t-statistic. Contours of the SMA and primary motor cortex are shown in green and blue respectively. (A) Upper panel displays regions where there was a significant interaction, such that there was greater cortical coherence with the STN than with the GPi at high rather than at low beta frequencies. The cluster encompasses mesial primary motor cortex and the SMA. Cross-hairs are centred on the location of the peak t-statistic at 8 - 46 82. In the lower panel the cross-hairs are centred at the location of the hand area of primary motor cortex, highlighting that the interaction effect was centred medially. (B) A cluster exhibiting a significant simple main effect of band for the GPi. This region encompasses lateral primary motor cortex and has greater coherence with the GPi at low rather than at high beta frequencies. Cross-hairs are centred on the location of the peak t-statistic at 38 -24 72. (C) Two clusters displaying a significant simple main effect of band for the STN. The clusters include SMA and mesial primary motor cortex. In the upper panel the cross-hairs are centred on the location of the peak t-statistic within the posterior cluster at MNI co-ordinates 8 -6 82. In the lower panel cross-hairs are centred on the location of the peak t-statistic within the anterior cluster at MNI co-ordinates 2 -48 58. (D), (E) and (F) show group mean coherence spectra (with standard errors) computed between the STN/GPi and cortical locations of the peak t-statistic of the simple main effect of band separately for the STN (corresponding to the anterior and posterior clusters in (C)) and GPi (corresponding to cluster in (B)). Black arrows indicate peaks within the low and high beta frequency ranges in STN coupling with cortex.

Additionally, we also observed reciprocal simple main effects of frequency band for the STN and GPi. **Figure 3B** indicates a focal lateral motor region in the vicinity of the hand area of primary motor cortex (peak *t* = 3.8, FWE p < 0.02 at MNI co-ordinates 38 -24 72) where cortico-GPi coherence was greater at low rather than at high beta frequencies. Conversely, **Figure 3C** depicts two mesial clusters that include SMA and the leg area of M1 (posterior cluster peak *t* = 4.5, FWE p = 3×10^−3^, at MNI co-ordinates 2 -48 58; anterior cluster peak *t* =4.2, FWE p = 6×10^−3^ at MNI co-ordinates 8 -6 82), where cortico-STN coupling is greater at high rather than at low beta frequencies.

To further visualise these effects, source extracted coherence spectra are shown in **Figures 3 D-F**. Source time series were extracted from the location of the peak *t* statistic within each cluster, for the simple main effect of band for each electrode location. Panels (D) and (E) show coherence profiles of the STN and GPi for the peak locations within the mesial anterior and mesial posterior clusters for which there was a simple main effect of band for the STN. Similarly, panel (F) shows the coherence profiles for the peak location within the single lateral cluster for which there was a simple main effect of band for the GPi. The cortical coherence profiles of the STN reveal distinct peaks within the low and high beta frequency ranges indicated by the black arrows in (D) and (E). Coupling with the GPi was intermediate, peaking between the black arrows and not extending as far to the right as for STN coupling with mesial motor areas.

### Directionality of beta band cortical-subcortical coupling and estimation of transmission delays

Based on the above evidence motor cortical coupling with the STN is divided into high and low beta frequencies. This is less pronounced in the profile of coupling between the same cortical areas and the GPi. We characterised the directionality of cortico-subcortical coupling at beta frequencies in order to determine whether these signals originate from the cortex or subcortical structures. For the purposes of this analysis we used source locations derived from the locations of the peak *t* statistics used for computation of the simple main effects as described above. Additionally, we divided the profile of cortico-STN coupling into high and low beta frequencies, but considered cortico-GPi coupling across the entire beta frequency band, given the lack of a clear distinction between beta sub-bands in coupling with GPi outside of that with a focal lateral motor region in the vicinity of the hand area of primary motor cortex.

For the STN, the difference in Granger causality was significantly greater than zero in the direction of cortex leading the STN for mesial but not for lateral motor regions, and at high but not low beta frequencies (**Figure 4A**). For the mesial anterior – STN network, one sample t tests were: *t* (24) = 3.96, *P* < 10^−3^ for the high beta sub-band and, and *t* (24) = 1.26, *P* > 0.1 for the low beta sub-band. Similarly, for the mesial posterior – STN network the statistics were as follows: *t* (24) = 2.21, *P* < 0.02 for the high beta sub-band and, and *t* (24) = −0.25, *P* > 0.5 for the low beta sub-band. Group level directionality within the lateral motor – STN network did not reach the threshold for significance: *t* (24) = 1.96, *P* > 0.05 for the high beta sub-band and, and *t* (24) = 1.12, *P* > 0.1 for the low beta sub-band. Similarly cortical activity in the studied mesial anterior and mesial posterior regions led activity within the GPi (**Figure 4B**): *t* (19) = 3.45, *P* < 2×10^−3^ for the mesial anterior region, *t* (19) = 2.33, *P* < 0.02 for the mesial posterior region and *t* (19) = 1.63, *P* > 0.05 for the lateral region.

**Fig 4.**
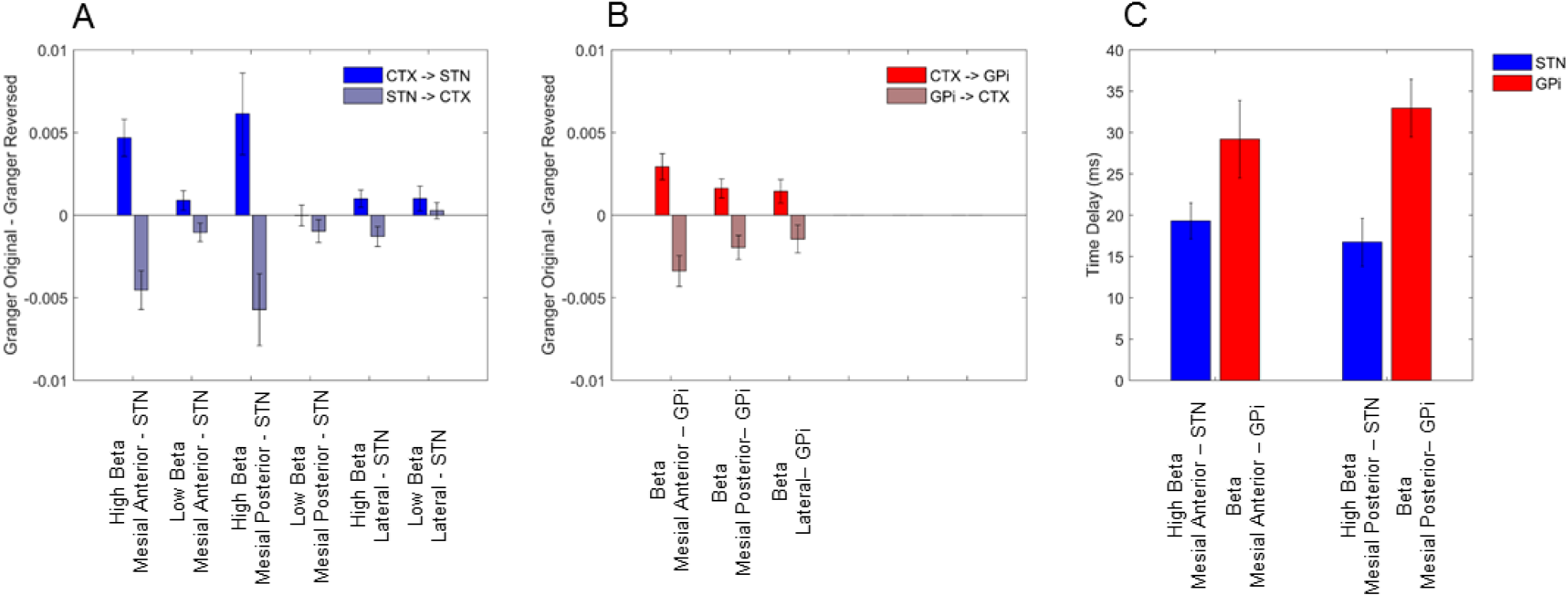
Granger directionality and analysis of net time delays: cortical driving of STN and GPi. Group mean differences in Granger causality between the original data and time reversed data are averaged across the high (21-30 Hz) and low (13-20 Hz) beta frequency ranges for the STN (A) and across the entire beta (13-30 Hz) frequency range for the GPi (B). Source time series for Granger causality computation are extracted from the locations of peak *t* statistics of the simple main effects of band, separately for the STN and the GPi as per Figure 3B and C. The difference in Granger causality is significantly greater than zero in the direction of mesial (anterior and posterior) motor cortical areas to the STN for the upper beta frequency band. Additionally, coupling in the direction of mesial and lateral motor areas to the GPi was also significant. For the same cortical areas that drive sub-cortical activity, Granger causality estimates are negative in the (reverse) direction of subcortical sites driving cortex, confirming that cortical activity leads the former. In the case of there being statistically significant unidirectional coupling, time delays between cortex and the STN/GPi were estimated (C). Time delays were estimated for the same cortical locations and frequency bands used for Granger causality analysis. Standard error bars are shown.

After directionality analysis, net time delays between mesial cortical regions and the STN/GPi were estimated (**Figure 4C**). Time delays were only estimated for these networks as there was predominant unidirectional coupling in the direction of mesial anterior and mesial posterior cortex to both the GPi and the STN. Furthermore, given that the GPi lies further downstream of the STN in the motor cortico-basal ganglia circuit, and lacks direct hyperdirect pathway inputs, we tested whether time delays from cortex to STN were shorter than those from cortex to GPi. Accordingly, we set up a 2 × 2 ANOVA to further investigate delays with factors cortical location (Mesial anterior vs Mesial posterior) and target location (STN vs GPi). Our results revealed a significant main effect of target location such that delays between mesial cortical areas and the GPi were longer than those between the same cortical areas and the STN (*F*(1,32) =15.81, *P* < 1×10^−3^). Importantly there was no significant main effect of cortical location (*F*(1,32) =0.02, *P* = 0.9), nor was there a significant interaction of the two factors (*F*(1,32) =1.55, *P* > 0.2).

Additionally, we determined whether the sharpness of the beta waveform, thought to reflect tight synchronisation to cortical inputs, might also provide support for dominance of a direct pathway from cortex to STN (see **Supplementary Results**).

### Functional connectivity is predicted by anatomical connectivity within the cortico-STN hyperdirect pathway

Based on our core hypothesis that coupling at high beta frequencies reflects activity within the hyperdirect pathway, we tested for a voxel wise relationship between cortico-STN tract density and cortico-STN high beta band coherence across all studied contacts for each hemisphere and each subject. The results of cluster based permutation testing are shown in **Figure 5A** and reveal a lateralised cluster encompassing the SMA, where tract density was predictive of high beta band coherence. The blue contour indicates the region where there was a significant simple main effect of band for the STN (this was the region where cortico-STN coherence was greater in the high beta band than in the low beta band; see **Figure 3C**), whilst the red and green contours indicate regions where at the group level high beta band coherence and tract density respectively, were greater than the 95% percentile. Further analysis revealed no significant relationship between cortico-STN tract density and cortico-STN low beta band coherence. Similar correlation of cortico-GPi tract density with cortico-GPi coherence in the high and low beta frequency bands was also not significant.

**Fig 5.**
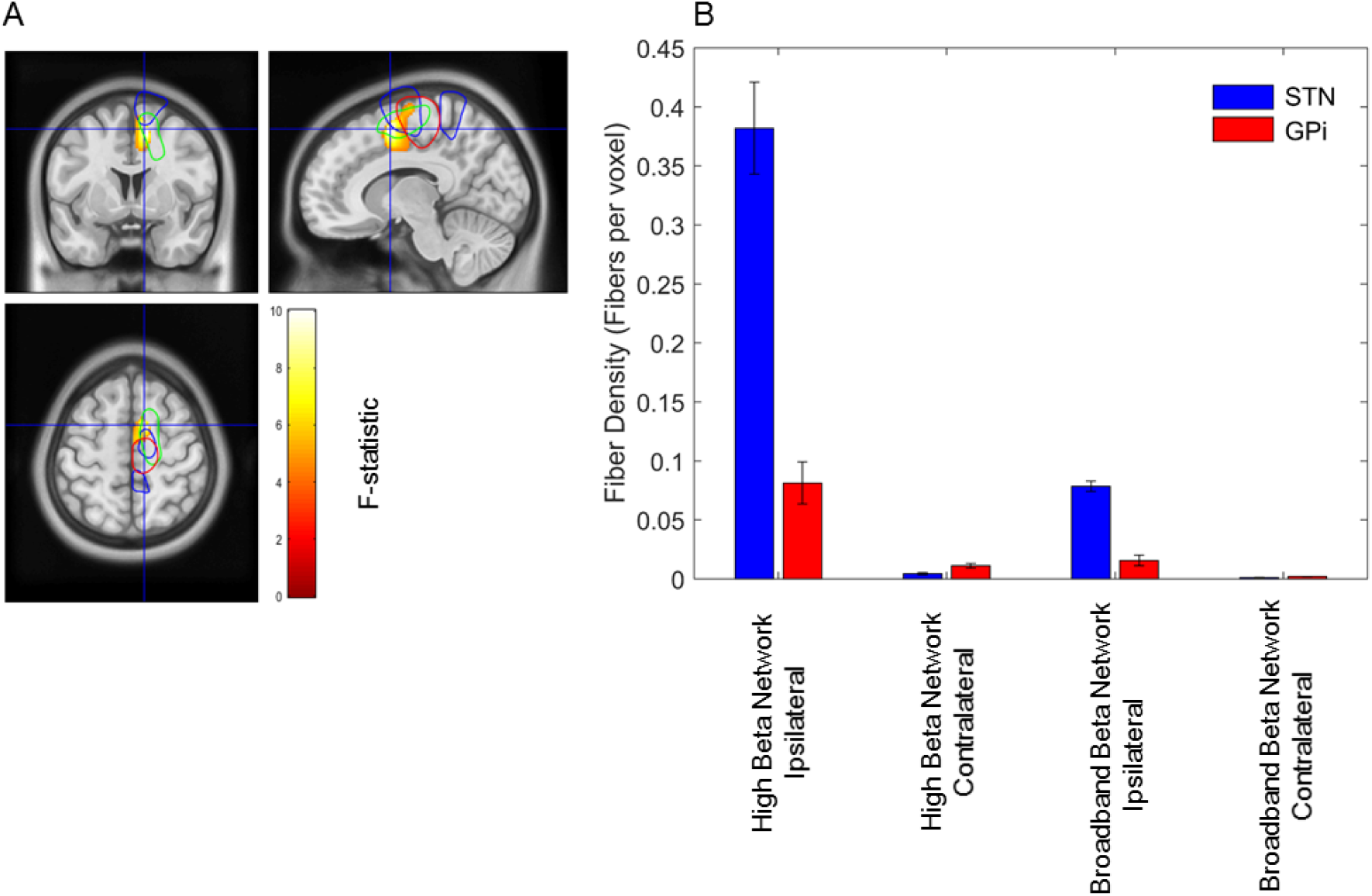
Relationship between structural and functional connectivity: tract density predicts hyperdirect pathway high beta coherence. (A): A statistical image of the significant cluster for the group level voxel wise correlation between high beta band cortico-STN coherence and cortico-STN fibre density. The image is superimposed on a T1-weighted MRI and the cross hairs are centred on the location of the peak F-statistic which lies within the SMA. The blue contour encloses the cortical volume where there was a simple main effect of band for the STN (see upper panel of Figure 3C). The red and green contours indicate regions where at the group level high beta band coherence and tract density respectively, were greater than the 95% percentile. (B): mean tract density estimates are plotted for the ‘high beta’ and ‘broadband beta’ networks separately for STN (blue) and GPi (red) contacts. Vertical bars represent standard errors of the mean.

For further visualisation of the overlap between MEG derived functional connectivity and tractography derived structural connectivity, we studied fibre tracts passing through the predefined spherical ROI for each contact which originated within cortical volumes for which: 1) there was a simple main effect of band for the STN (the areas for which cortico-STN coherence was greater at high rather than at low beta frequencies – which we name the ‘high beta network’) and 2) coherence with the STN/GPi across the entire beta frequency range (which we name the ‘broadband beta network’) was greater than coherence at alpha band frequencies. Group analyses are displayed in **Supplementary Figure 5**.

Tract densities were computed for the ipsilateral and contralateral high beta and broadband beta networks, separately for STN and GPi contacts. Results are summarised in **Figure 5B** and reveal that STN contacts have relatively dense fibre innervations from ipsilateral cortical regions that couple to them preferentially at high beta frequencies. We explored this phenomenon statistically by constructing a 2 × 2 × 2 factorial ANOVA with factors network (high beta vs. beta), laterality (ipsilateral vs. contralateral) and electrode location (STN vs. GPi). Significant main effects were observed for all three factors (high beta vs. low beta: *F*(1,215) =44.16, *P* < 10^−4^; ipsilateral vs. contralateral: *F*(1,215) =136.37, *P* < 10^−4^; STN vs. GPi: *F*(1,215) =54.03, *P* < 10^−4^). Additionally, all pairwise interactions and the three way interaction were significant, (interaction of electrode location and laterality: *F*(1,215) = 45.46, *P* < 10^−4^; interaction of electrode location and network: *F*(1,215) = 25.77, *P* < 10^−4^; interaction of network and laterality *F*(1,215) = 50.18, *P* < 10^−4^; interaction of network, laterality and electrode location *F*(1,215) = 25.70, *P* < 10^−4^), highlighting that the difference in tract density between the high beta and broadband beta networks was greater for the ipsilateral rather than the contralateral STN, and that this effect was less marked for the GPi.

### Computational model describing differential mechanisms by which high and low beta frequencies are generated in the cortico-basal-ganglia circuit

We have provided empirical evidence that the hyperdirect pathway drives high beta activity in the basal ganglia. However, what is the origin of beta activity in the lower frequency band and to what extent are the two activities functionally distinct ^4,31,32^? It is after all the low beta activity that is believed to be pathological. To answer these questions, we turned to computational modelling and designed a model able to capture a number of empirical features observed both in our own data, and in those of other electrophysiological studies. Work in computational neuroscience suggests that circuits composed from homogeneous populations of excitatory and inhibitory neurons may generate oscillations with single dominant frequency ^33^. To produce two frequencies in the model, we included two pairs of excitatory-inhibitory network generators: 1) an excitatory-inhibitory cortical network capable of generating high beta frequency oscillations and 2) the reciprocal excitatory-inhibitory STN-GPe subcortical network which served to generate low beta frequency oscillations ^18,34^.

The structure of our model and its ability to generate high beta activity cortically and low beta activity subcortically is illustrated in **Supplementary Figure 6**.

### Modelling the effects of cortico-subcortical connectivity on the frequency and coherence of oscillations

**Figure 6** illustrates the effects of cortical input on the frequencies of oscillations within the STN-GPe loop. The left image in (A) shows the effect of modulating the hyperdirect pathway connection strength, W_ES_ on the simulated spectra of the STN and GPi. As W_ES_ is increased from 0, high beta oscillations from cortex start to appear subcortically. At values of W_ES_ between 5 and 10, low beta frequency oscillations (~13 Hz) appear in the spectra of STN and GPi and then disappear when W_ES_ exceeds a value of 10. The middle and right most image in (A) explore the effects of varying cortical time constants (*τ* _*E*_/*τ* _*I*_) and the delay between the two cortical populations (T_EI/IE_). Varying these cortical parameters primarily influences the frequency of the high beta oscillatory activity incoming from the cortex, with little effect on the low beta peak frequency.

**Fig 6.**
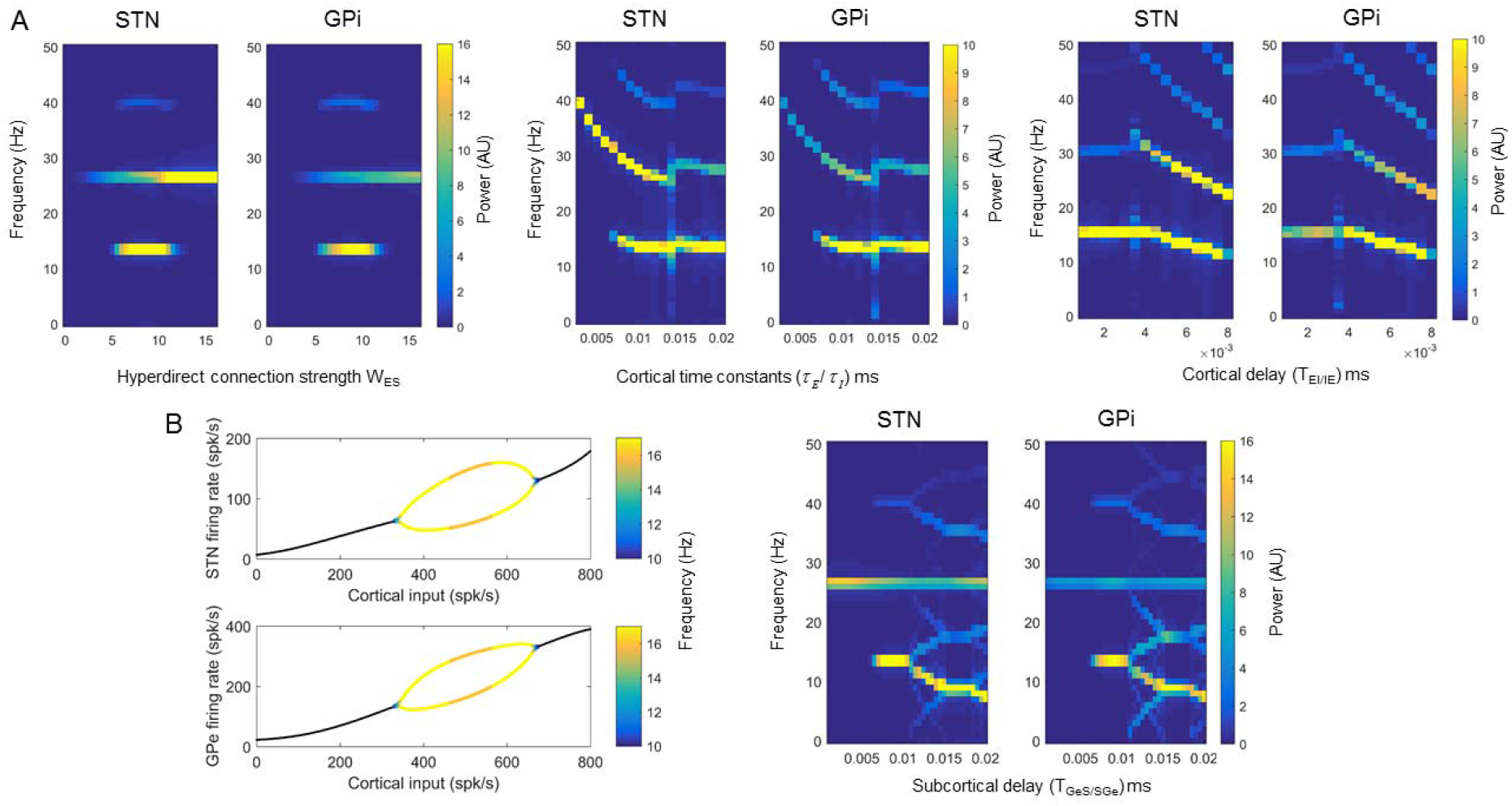
Computational modelling: the cortex generates high beta activity which is propagated subcortically and provokes the generation of pathological lower beta frequencies. (A): We explore the effect of altering the hyperdirect connection strength (W_ES,_ left figure), cortical time constants (*τ* _*E*_/*τ* _*I*_, middle figure) and the cortical delay (T_IE_/T_IE_, right figure) on the spectra of the STN and GPi. High beta frequency activity generated within the cortex propagates to the STN and GPi. Additionally under certain parameter values, a lower beta frequency (between 10-15 Hz) emerges. The left image in (B) is a bifurcation diagram highlighting that low beta frequency oscillatory activity may be generated in the STN-GPe feedback loop depending on cortical inputs. In the oscillatory region of STN and GPe firing the dominant frequency is colour coded. The right image in (B) displays the effect on STN and GPi spectra of changing the subcortical delay parameters (T_GeS_ /T_SGe_).

Our simulations indicate how a strong hyperdirect pathway may lead to the generation of low beta frequency oscillations subcortically. To illustrate this more clearly, in the left image of **Figure 6B** is a bifurcation diagram for STN and GPe activities for a reduced model of only the reciprocal STN-GPe network, where both populations receive striatal input and the STN receives fixed (non-oscillatory) excitatory cortical inputs. The x-axis plots the cortical input which is the product of the hyperdirect pathway connection strength, W_ES_ and the firing rate of the cortical excitatory population. The system displays a stable fixed point (black lines) which transitions to an instability and oscillatory behaviour at cortical input values between approximately 330-670 spikes/second. Within the oscillatory range the peak of the low beta frequency is indicated in the colour bar. Interestingly this simulation predicts that very high hyperdirect pathway strengths lead to the loss of low beta oscillations in the sub-cortex. The right image in **Figure 6B** shows how the peak frequency of the low beta oscillation generated in the STN-GPe loop can be influenced by the transmission delays between these two structures (T_GeS/SGe_).

Next, using our model we simulated power spectra of the cortex, STN and GPi and coherences between cortex-STN and cortex-GPi (**Figure 7A)**. Parameters used are listed in **Supplementary Table 2,** but we varied the value of the strength of the net inhibitory loop, W_GiE,_ between the GPi, thalamus and cortex. In these simulations noise was added (see **Supplementary Methods**) in order to make coherence values physiologically plausible. For a range of values of W_GIE_ oscillatory peaks within the high and low beta frequency ranges are seen in the STN and GPi. As observed in our own data (**Figures 2 and 3**), both high beta band power and cortical coherence were greater for the STN than for the GPi. Increasing W_GiE_ led to an increase in the amplitude of the low beta frequency peak in both the STN and the GPi in addition to increases in cortico-STN and cortico-GPi coherence at low beta frequencies. Interestingly, cortical activity predominantly displays a high beta frequency peak without the low beta frequency peak observed in subcortical activity. This is in keeping with electrocorticographic studies in PD patients which display spectral peaks above 20 Hz ^35–37^.

**Fig 7.**
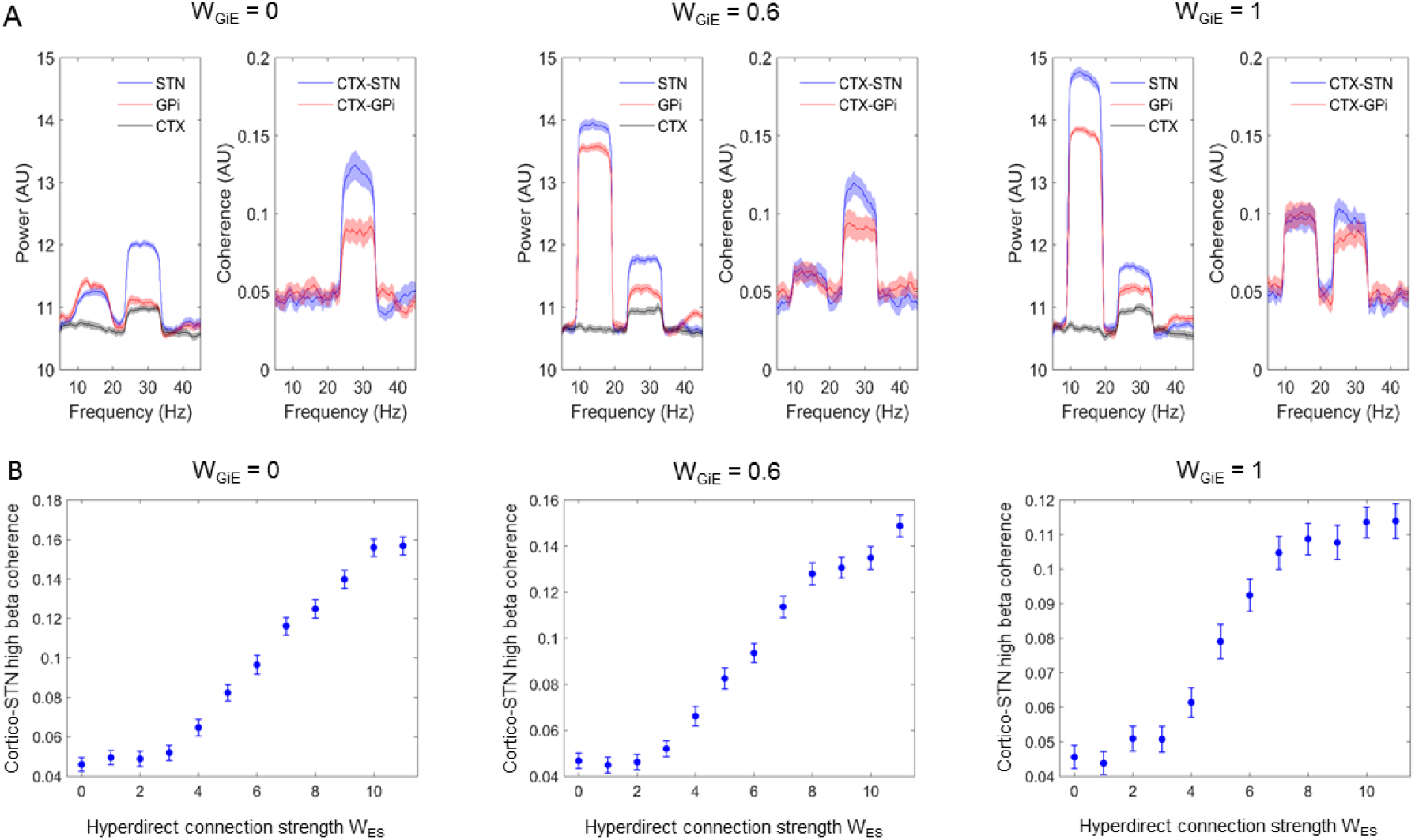
Computational modelling captures observed power and coherence spectra and predicts the observed relationship between hyperdirect pathway structural and functional connectivity. (A): Simulated power spectra for the STN, GPi and cortex are shown with the associated profiles of cortico-STN and cortico-GPi coherence spectra. Parameter values for this simulation are as per **Supplementary Table 2**, except for the fact that the strength of the net inhibitory loop, W_GIE,_ between the GPi, thalamus and cortex was varied as shown. Power and coherence spectra display peaks at high and low beta frequencies, in keeping with experimental data. Increasing W_GIE_ results in increased low beta frequency power and coherence. (B): the simulated relationship between hyperdirect pathway strength and cortico-STN high beta band coherence displays a monotonically increasing profile. Standard errors of the mean are computed over 50 simulations.

Finally **Figure 7B** reveals that the computational model developed predicts a monotonically increasing relationship between hyperdirect pathway strength W_ES_ and cortico-STN coherence in the high beta frequency range (21-30 Hz). The slope of this relationship is greatest above values of W_ES_ that lead to the propagation of high beta oscillations sub-cortically (see **Figure 6A**).

In summary, modelling reveals that an exaggerated hyperdirect pathway may be a pre-requisite for the generation of subcortical low beta frequency rhythms, and that connections from basal ganglia via thalamus to cortex may underlie the coherence between GPi/STN and cortex at low beta frequencies. The model also captures the empirical relationship between structural and functional connectivity within the hyperdirect pathway.

## Discussion

This study characterised the intersection between structural and functional connectivity within the basal ganglia in PD patients undergoing DBS surgery. The hyperdirect pathway from cerebral cortex almost exclusively terminates in the STN, and so by comparing LFPs from patients with STN electrodes to those from patients with GPi electrodes, we were able to isolate electrophysiological markers of hyperdirect pathway activity. STN activity displayed both greater local synchrony, as indexed by LFP amplitude, and greater functional connectivity with motor cortical areas, including the SMA and mesial primary motor cortex, at high beta frequencies. Furthermore, these same cortical regions tended to drive STN activity within the high beta frequency range with shorter delays than those that were observed for coupling to the GPi. The extrema of beta oscillations were also sharper in the STN than in the GPi after matching for both frequency and power, in keeping with the notion of a more synchronised hyperdirect cortical input to the former. More strikingly we show across participants that cortico-STN coherence at high beta frequencies correlates with cortico-STN fibre tract densities in a focal premotor region encompassing the SMA – an area that plays a key role in the volitional control of movement ^38^. Collectively these data provide evidence that hyperdirect pathway activity within the cortico-basal-ganglia circuit evokes a unique spectral signature at high beta frequencies.

We interpret the data in the context of a computational model which provides novel insight into the origins of beta oscillatory activity in PD. Our model demonstrates how an exaggerated hyperdirect pathway in PD is capable of both generating high beta frequency oscillations and inducing synchrony at lower beta frequencies within subcortical circuits. The model therefore provides a formal biophysical mechanism for frequency transduction in cortico-basal ganglia circuits, which hitherto has only been speculated ^4,32^. Excessive sub-cortical synchrony at low beta frequencies is considered to be closely related to motor impairment in PD ^39^ and therefore strategies aimed at modulating the hyperdirect pathway may exert therapeutic effect by reducing subcortical transduction to pathological low beta activity.

### Intersection between structural and functional connectivity within the cortico-basal-ganglia circuit

We report a novel correlation between cortico-STN coherence and cortico-STN fibre tract density that was specific anatomically to mesial premotor areas (SMA) and spectrally to upper beta band frequencies. Importantly this correlation was not observed in the profile of cortical coherence with the GPi. We interpret this finding as being indicative of a tight relationship between structural and functional connectivity within the monosynaptic hyperdirect pathway from the cortex to the STN ^9,15,40^. Of particular relevance to our findings is the strong theoretical relationship between functional and effective – and implicitly also structural - connectivity in networks such as the hyperdirect pathway with unidirectional information transfer ^41^.

Previous studies examining the intersection of tractography and fMRI are suggestive of a similar overlap of structural and functional connectivity within hyperdirect connections during stopping behaviour ^10,42^ with the strength of hyperdirect connections also correlating with the efficacy of stopping ^43^. More broadly, the relationship between functional and structural connectivity in brain networks is highly complex ^44^ and also limited by the pitfalls of the modalities used to assess these measures. Nevertheless, significant overlaps between structural and functional connectivity have been noted in highly conserved networks such as the default mode network ^45^.

We hypothesise that the lack of an observed correlation between structural and functional connectivity within the cortico-GPi network is indicative of the fact that the GPi receives less synchronised cortical inputs through spatiotemporal intermixing of hyperdirect, direct and indirect pathway inputs that are themselves mediated by other nuclei before reaching the GPi ^46^. Interestingly however the existence of a direct connection between the cortex and the GPi has recently been postulated and this would explain our finding of fibres passing between these two structures which bypass the striatum ^47^. Our data suggest that this pathway is likely to be less dominant than the hyperdirect pathway based on the comparison of tract densities.

Finally, we estimated a relatively long delay (>15ms) for cortico-STN hyperdirect transmission in the high beta band. This may arise because activities in the hyperdirect and indirect pathways are only relatively and not absolutely spectrally distinct at the level of the STN. Therefore estimates of delay in the hyperdirect pathway will be exaggerated by the indirect component overlapping in frequency ^4,48^. Estimated delays also not only include conduction delays but also delays for synaptic integration, which increase as more synapses are involved, and may be extended by inhibitory inputs to the STN.

### Oscillatory non-sinusoidality as a marker of synchronised cortical inputs

STN and GPi beta activities were not only distinguished by relatively different peak frequencies, and patterns of cortical coupling and corresponding delays, but also by the shape taken by the oscillations themselves. The extrema of beta oscillations were sharper in the STN LFP than the GPi LFP, even when beta power and frequencies were matched. The sharpness of oscillatory population activities has increasingly been noted to have functional significance. For example, the sharpness of cortical beta oscillations in PD correlates with rigidity ^30^, and is reduced during DBS ^30^ and after dopaminergic medication ^49,50^. Sharpness is thought to be associated with tight synchronisation in rhythmic inputs, although the evidence supporting this interpretation has so far been theoretical rather than empirical in nature ^29,30,51–53^. Still this interpretation would explain the greater sharpness of oscillations in the STN where the mono-synaptic hyperdirect pathway is relatively dominant, and is supported by the striking correlation between beta oscillation sharpness and hyperdirect tract density amongst the subthalamic LFP recordings.

### Insights from the computational modelling

The model presented here underscores the importance of an exaggerated hyperdirect pathway in PD, which has also been suggested in previous reports ^4,18,21,40^. More specifically we show for the first time how the exaggerated subcortical propagation of high beta frequencies via the hyperdirect pathway can lead to the generation of the lower beta frequencies that are believed to play a more direct role in the genesis of motor impairment ^39^. This explanation is consistent with our own and previous findings that the net directionality of high beta frequency oscillations is from cortex to STN ^4,54^. In contrast we observed no significant net directionality for low beta frequency oscillations, which may reflect low subcortico-cortical feedback strengths via the GPi-thalamo-cortical loop (parameter W_GiE_ in **Supplementary Table 2**). Secondly, our model predicted a monotonically increasing relationship between hyperdirect pathway strength and cortico-STN high beta band coherence, which is in keeping with the observed tight relationship between *structural* and *functional* connectivity within the hyperdirect pathway. The model also captures the relative difference in local and cortical synchrony in the high beta band for the STN and GPi, in the context of strong hyperdirect pathway inputs.

Further in keeping with our model is the finding that DBS - which may exert therapeutic benefit by suppressing the hyperdirect pathway ^4,14,21,55^ – has the effect of reducing coherence in the upper beta frequency band between the cortex and the STN and also the cortex and the GPi, in addition to suppressing low beta synchrony locally within the STN and the GPi ^4,37,56^.

### Study Limitations

Our findings should be considered in light of the following limitations. Firstly, we observed phenotypic differences between the STN and GPi DBS patient groups. Patients implanted in the GPi suffered from greater motor and cognitive impairment and there was a tendency for them to receive lower levodopa equivalent doses than patients implanted in the STN. Of these factors, both elevated motor impairment and reduced levodopa doses would tend to increase LFP beta power in the GPi subgroup compared to the STN subgroup ^22,57,58^, therefore potentially leading to an underestimation of the group differences observed. We accounted for phenotypic differences in statistical analyses by including motor and cognitive scores and levodopa equivalent doses as covariates.

We note that similar phenotypic differences are reported in PD patients undergoing DBS of the STN or GPi, and that such differences reflect predominant symptomatology guiding target selection ^59^. It is often argued that the GPi may be a more appropriate target than the STN in patients with existing mild cognitive dysfunction or troublesome dyskinesias ^60,61^. The latter may necessitate reduced medication doses and consequently increased on medication UPDRS III scores, as was the case in our cohort.

Secondly, patients were recorded on medication. Whilst this is likely to have reduced the low beta power and coherence seen in our recordings, it is unlikely to have significantly impacted high beta band power in the STN or GPi, or indeed coherence between cortex and these structures ^2,31,57,62^.

Finally, we utilised connectome data from a large cohort of patients with Parkinson’s disease rather than connectivity data from individual patients. Although this may be considered a limitation, a potential advantage is the fact that connectome datasets are acquired using special MRI hardware with large cohort sizes leading to connectivity estimates with improved signal-to-noise ratio compared to what would be possible with individual patient data. Connectome data have been successfully leveraged to study the mechanisms of DBS action ^63^ and also to explore how data from disparate lesion studies can be integrated to understand the role of brain networks in disease ^64^.

## Materials and Methods

### Patients and experimental details

Twelve patients; six with bilateral implantation of STN DBS electrodes and six with bilateral implantation of GPi DBS electrodes were recruited. Patients were diagnosed with Parkinson’s disease according to the Queen Square Brain Bank criteria ^65^. Patients in the GPi DBS subgroup had an additional diagnosis of Parkinson’s disease dementia (PDD) and were recruited into a separate trial which involved targeting both the GPi and the Nucleus Basalis of Meynert (NBM) ^66^. Clinical characteristics of the patients are presented in **Supplementary Table 1**. The STN and GPi DBS subgroups did not differ in age or disease duration (two-tailed, unpaired t tests for: 1) age, *t* = −1.6, *p* = 0.13 and 2) disease duration, *t* = −0.92, *p* = 0.38). There was a trend towards the levodopa equivalent dose being higher in the STN subgroup (*t* = 1.9, *p* = 0.084). This group also had a lower UPDRS Part III motor score on medication and higher cognitive performance scores as measured by the Mini-Mental State Examination (for UPDRS Part III motor score, *t* = −2.87, *p* = 0.02; for MMSE, *t* = 7.6, *p* < *p* < 10^−4^). In order to account for phenotypic differences, we included the three clinical features which were significantly different between the groups as covariates in statistical analyses. 275-channel MEG data was collected simultaneously with LFP activity recorded from DBS electrodes sited in the STN or GPi. Recordings were performed on medication and at rest. Further details of the operative procedure and recordings are found in the **Supplementary Methods**.

### DBS electrode localisation and fibre tracking

DBS electrodes were localised using Lead-DBS (www.lead-dbs.org). This involved linearly co-registering the post-operative MRI to the pre-operative MRI using SPM12 (Statistical Parametric Mapping; http://www.fil.ion.ucl.ac.uk/spm/software/spm12). In order to then compare electrode placement across subjects, pre- and postoperative MRI acquisitions were non-linearly co-registered (normalisation) into MNI ICBM152 NLIN 2009b stereotactic space using the SPM12 segment nonlinear option in Lead-DBS. Each electrode could then be localised and visualised in the aforementioned MNI space simultaneously with masks of subcortical structures (including GPi and STN) derived from the DISTAL atlas within Lead-DBS.

For each electrode we used an automated approach for determining which individual contacts lay within the STN or GPi. This approach relied on determining whether MNI co-ordinates defining individual contacts lay within a convex hull bounded by the surface of the STN or GPi (https://uk.mathworks.com/matlabcentral/fileexchange/10226-inhull). Contact pairs where at least one contact lay in the target nucleus were used for subsequent analyses; for instance, in the event of only contact 1 being inside the target nucleus, we selected contact pairs 0-1 and 1-2. For the purposes of fibre tracking, a spherical region of interest centred at the midpoint of each chosen contact pair, with a radius that just encompassed each contact pair was constructed. This spherical volume was used as a seed region in an openly available group connectome (www.lead-dbs.org) which was derived from diffusion-weighted magnetic resonance images of 90 patients in the Parkinson’s progression markers initiative (PPMI) database. All scanning parameters are published on the website (www.ppmi-info.org). Whole brain tractography fibre sets were calculated using a generalized q-sampling imaging algorithm as implemented in DSI studio (http://dsi-studio.labsolver.org) within a white-matter mask after segmentation with SPM12. Fibre tracts were transformed into MNI space ^67^ for visualisation. We then determined the number of fibres passing through both the aforementioned spherical seed and each cubic voxel of side 2mm. This yielded a single number for each voxel that served as an estimate of tract density and was written to a 3D image. Prior to statistical testing images were smoothed with an 8mm isotropic Gaussian kernel as per the analysis of MEG data.

### Analysis of oscillatory synchrony within the cortico-STN/cortico-GPi circuit

Power spectra from STN and GPi contact pairs were computed using multitaper spectral estimation with a frequency resolution and taper smoothing frequency of 2.5 Hz ^68^. Physiological power spectra may be thought of as a summation of two distinct processes: 1) an aperiodic component reflecting 1/f like characteristics which may differ across subjects and 2) periodic oscillatory components manifesting as band limited peaks in the power spectrum. To make spectra comparable across subjects we used a spectral parameterisation algorithm (Fitting Oscillations & One-Over F algorithm, https://github.com/fooof-tools/fooof) to model the aperiodic (1/f) component. This was subsequently subtracted from the power spectrum in order to isolate the periodic oscillatory component of interest. To test for differences in the spectra of STN and GPi at each frequency we used an independent samples t-test, with correction for multiple comparisons using cluster based permutation testing (P<0.01 corrected, cluster forming threshold P<0.01).

Brain areas coherent with STN and GPi LFPs were localized using dynamic imaging of coherent sources (DICS) beamforming ^69^ yielding a 3D image of coherence (see **Supplementary Methods** for further details). In light of previous work highlighting differences in the profile of STN-cortical connectivity in the upper (21-30 Hz) and lower (13-21 Hz) frequency bands, we restricted our DICS beamformer analysis to these two bands ^4^. For each subject and each hemisphere, coherence images were generated for the two frequency bands. Half of the resulting images (all left STN/GPi images) were reflected across the median sagittal plane to allow comparison of ipsilateral and contralateral sources regardless of original side. These images were then subjected to a 2 × 2 factorial ANOVA, with frequency (low beta versus high beta) and electrode location (STN versus GPi) as factors in SPM.

In addition to modelling subject-specific dependencies in the recordings from the two hemispheres, we included side as an additional categorical variable for each subject to account for potential differences between the recordings from the right and left sides. Covariates representing each patient’s preoperative levodopa equivalent dose, UPDRS Part III motor score on medication and MMSE were introduced as described above in order to account for phenotypic differences between the two groups. We tested for the direction of main effects and interactions in the 2 × 2 ANOVA by performing t-tests in SPM. All analyses were corrected for multiple comparisons using random field theory and reported findings are significant with familywise error (FWE) correction at the cluster level (*P* < 0.01 corrected, cluster forming threshold *P* < 0.001 uncorrected). DICS beamformer images were also generated across the alpha (7-13 Hz) and entire beta (13-30 Hz) frequency and subjected to a separate 2 × 2 factorial ANOVA with factors frequency (alpha versus beta) and electrode location (STN versus GPi). As the aim of this analysis was to define a cortical network coupled to the STN/GPi across the entire beta frequency range - for subsequent visualisation of tracts (see Results) - we were primarily interested in the main effect of band.

Next, using beamforming we performed time series extraction from peak voxels in the SPMs of group level main effects and interactions. Coherence was computed between the reconstructed source and the subcortical LFP using multitaper spectral estimation with a frequency resolution and taper smoothing frequency of 2.5 Hz ^68^. Further details of time series analysis relating to directionality and waveform shape are found in the **Supplementary Methods**.

### Relationship of structural and functional connectivity

For each contact pair (with at least one contact lying in the STN or GPi) in each subject, we had isotropic 2mm 3D images of both coherence and fibre density in MNI space. Further analysis was performed separately for each patient group (STN electrodes versus GPi electrodes) and each frequency band (low beta 13-21 Hz, and high beta 21-30 Hz) in order to establish whether tract density was predictive of coherence. As we were specifically interested in fibres representing hyperdirect connections, we minimised the chance of including fibres from the direct and indirect pathways by limiting fibre tracts to those not traversing the striatum.

For each voxel we constructed a General Linear Model (using spm_ancova) with tract density as the independent variable and coherence as the dependent variable. Subject, side, medication dose and MMSE were introduced as covariates as described above. The F-statistic was used to determine a P value, with a threshold of P<0.01 being used to define voxels to form clusters for cluster based permutation testing. This served to correct for multiple comparisons and we utilised a threshold of P<0.01 to define significant clusters. The F-statistics of voxels within each significant cluster were then written to a 3D image for visualisation. In further analysis we investigated fibre tracts traversing the spherical ROI associated with each contact, which started (or terminated) within cortical volumes derived from SPM analysis of main effects of cortico-STN and cortico-GPi coherence. For each contact we computed a single overall tract density estimate by dividing the number of fibres originating in each cortical volume by the number of voxels contained within the volume.

### Computational modelling of high and low beta band oscillatory synchrony

A firing rate model of the cortico-basal-ganglia circuit was developed, based on models previously used to study beta oscillations ^18,34^. The basic idea behind our model is to generate oscillations via the interaction of excitatory and inhibitory neural populations. Technical details are described in **Supplementary Methods**.

## Author contributions

Study design and methodology, AO, VL, RB, MH and PB. Contribution of software, AO, CY, WJN, AH, VL. MEG and LFP data collection AO, JG, VL. Data analysis, AO and CY. Clinical data collection and patient characterisation, AO, HA, LZ, TF, PL. Writing – original draft AO and PB. Writing – review & editing AO, CY, WJN, JG, HA, AH, RB, MH, PB, VL. Funding acquisition AO, VL and PB. Supervision VL and PB.

## Data and materials availability

The MEG and LFP datasets generated within the current study have not been deposited within a public repository because they contain patient sensitive data. Anonymised datasets are available from Dr Ashwini Oswal on reasonable request however. MATLAB code for reproducing: 1) the relationship between structural and functional connectivity, 2) computation of time delays and 3) results of computational modelling are deposited on the GitHub repository https://github.com/AshOswal/Multimodal_Tools.

## References

1. Hirschmann, J. et al. Distinct oscillatory STN-cortical loops revealed by simultaneous MEG and local field potential recordings in patients with Parkinson’s disease. Neuroimage 55, 1159–68 (2011).

2. Litvak, V. et al. Resting oscillatory cortico-subthalamic connectivity in patients with Parkinson’s disease. Brain 134, 359–74 (2011).

3. Williams, D. et al. Dopamine-dependent changes in the functional connectivity between basal ganglia and cerebral cortex in humans. Brain 125, 1558–69 (2002).

4. Oswal, A. et al. Deep brain stimulation modulates synchrony within spatially and spectrally distinct resting state networks in Parkinson’s disease. Brain 139, 1482–1496 (2016).

5. Albin, R. L., Young, A. B. & Penney, J. B. The functional anatomy of basal ganglia disorders. Trends Neurosci. 12, 366–375 (1989).

6. DeLong, M. R. & Wichmann, T. Circuits and circuit disorders of the basal ganglia. Archives of Neurology 64, 20–24 (2007).

7. DeLong, M. R. Primate models of movement disorders of basal ganglia origin. Trends Neurosci. 13, 281–5 (1990).

8. DeLong, M. & Wichmann, T. Update on models of basal ganglia function and dysfunction. Park. Relat. Disord. 15, (2009).

9. Nambu, A., Tokuno, H. & Takada, M. Functional significance of the cortico-subthalamo-pallidal ‘hyperdirect’ pathway. Neurosci. Res. 43, 111–7 (2002).

10. Aron, A. R., Behrens, T. E., Smith, S., Frank, M. J. & Poldrack, R. A. Triangulating a cognitive control network using diffusion-weighted Magnetic Resonance Imaging (MRI) and functional MRI. J. Neurosci. 27, 3743–3752 (2007).

11. Chen, W. et al. Prefrontal-subthalamic hyperdirect pathway modulates movement inhibition in humans. Neuron 1–10 (2020). doi:10.1016/j.neuron.2020.02.012

12. Haynes, W. I. A. & Haber, S. N. The organization of prefrontal-subthalamic inputs in primates provides an anatomical substrate for both functional specificity and integration: Implications for basal ganglia models and deep brain stimulation. J. Neurosci. 33, 4804–4814 (2013).

13. Ashby, P. et al. Potentials recorded at the scalp by stimulation near the human subthalamic nucleus. Clin. Neurophysiol. 112, 431–7 (2001).

14. Miocinovic, S. et al. Cortical potentials evoked by subthalamic stimulation demonstrate a short latency hyperdirect pathway in humans. J. Neurosci. 38, 9129–9141 (2018).

15. Lambert, C. et al. Confirmation of functional zones within the human subthalamic nucleus: patterns of connectivity and sub-parcellation using diffusion weighted imaging. Neuroimage 60, 83–94 (2012).

16. Akram, H. et al. Subthalamic deep brain stimulation sweet spots and hyperdirect cortical connectivity in Parkinson’s disease. Neuroimage 158, 332–345 (2017).

17. Cagnan, H., Denison, T., McIntyre, C. & Brown, P. Emerging technologies for improved deep brain stimulation. Nature Biotechnology 37, 1024–1033 (2019).

18. Pavlides, A., Hogan, S. J. & Bogacz, R. Computational Models Describing Possible Mechanisms for Generation of Excessive Beta Oscillations in Parkinson’s Disease. PLoS Comput. Biol. 11, e1004609 (2015).

19. Moran, R. J. et al. Alterations in Brain Connectivity Underlying Beta Oscillations in Parkinsonism. PLoS Comput. Biol. 7, e1002124 (2011).

20. Marreiros, A. C., Cagnan, H., Moran, R. J., Friston, K. J. & Brown, P. Basal ganglia-cortical interactions in Parkinsonian patients. Neuroimage 66C, 301–310 (2012).

21. Gradinaru, V., Mogri, M., Thompson, K. R., Henderson, J. M. & Deisseroth, K. Optical deconstruction of parkinsonian neural circuitry. Science 324, 354–9 (2009).

22. Brown, P. et al. Dopamine dependency of oscillations between subthalamic nucleus and pallidum in Parkinson’s disease. J. Neurosci. 21, 1033–8 (2001).

23. Eusebio, A. et al. Deep brain stimulation can suppress pathological synchronisation in parkinsonian patients. J. Neurol. Neurosurg. Psychiatry 82, 569–73 (2011).

24. Whitmer, D. et al. High frequency deep brain stimulation attenuates subthalamic and cortical rhythms in Parkinson’s disease. Front. Hum. Neurosci. 6, 155 (2012).

25. Kühn, A. A., Kupsch, A., Schneider, G.-H. & Brown, P. Reduction in subthalamic 8-35 Hz oscillatory activity correlates with clinical improvement in Parkinson’s disease. Eur. J. Neurosci. 23, 1956–60 (2006).

26. Kühn, A. A. et al. Pathological synchronisation in the subthalamic nucleus of patients with Parkinson’s disease relates to both bradykinesia and rigidity. Exp. Neurol. 215, 380–7 (2009).

27. Weinberger, M. et al. Beta oscillatory activity in the subthalamic nucleus and its relation to dopaminergic response in Parkinson’s disease. J. Neurophysiol. 96, 3248–56 (2006).

28. Ray, N. J. et al. Local field potential beta activity in the subthalamic nucleus of patients with Parkinson’s disease is associated with improvements in bradykinesia after dopamine and deep brain stimulation. Exp. Neurol. 213, 108–13 (2008).

29. Cole, S. R. & Voytek, B. Brain Oscillations and the Importance of Waveform Shape. Trends in Cognitive Sciences 21, 137–149 (2017).

30. Cole, S. R. et al. Nonsinusoidal beta oscillations reflect cortical pathophysiology in parkinson’s disease. J. Neurosci. 37, 4830–4840 (2017).

31. López-Azcárate, J. et al. Coupling between beta and high-frequency activity in the human subthalamic nucleus may be a pathophysiological mechanism in Parkinson’s disease. J. Neurosci. 30, 6667–77 (2010).

32. Brittain, J. S. & Brown, P. Oscillations and the basal ganglia: Motor control and beyond. NeuroImage 85, 637–647 (2014).

33. Tiesinga, P. & Sejnowski, T. J. Cortical Enlightenment: Are Attentional Gamma Oscillations Driven by ING or PING? Neuron 63, 727–732 (2009).

34. Holgado, A. J. N., Terry, J. R. & Bogacz, R. Conditions for the generation of beta oscillations in the subthalamic nucleus-globus pallidus network. J. Neurosci. 30, 12340–52 (2010).

35. Crowell, A. L. et al. Oscillations in sensorimotor cortex in movement disorders: An electrocorticography study. Brain 135, 615–630 (2012).

36. De Hemptinne, C. et al. Therapeutic deep brain stimulation reduces cortical phase-amplitude coupling in Parkinson’s disease. Nat. Neurosci. 18, 779–786 (2015).

37. Wang, D. D. et al. Pallidal deep-brain stimulation disrupts pallidal beta oscillations and coherence with primary motor cortex in Parkinson’s disease. J. Neurosci. 38, 4556–4568 (2018).

38. Nachev, P., Kennard, C. & Husain, M. Functional role of the supplementary and pre-supplementary motor areas. Nature Reviews Neuroscience 9, 856–869 (2008).

39. Hammond, C., Bergman, H. & Brown, P. Pathological synchronization in Parkinson’s disease: networks, models and treatments. Trends Neurosci. 30, 357–64 (2007).

40. Baudrexel, S. et al. Resting state fMRI reveals increased subthalamic nucleus-motor cortex connectivity in Parkinson’s disease. Neuroimage 55, 1728–38 (2011).

41. Friston, K. J. et al. DCM for complex-valued data: Cross-spectra, coherence and phase-delays. Neuroimage 59, 439–455 (2012).

42. Aron, A. R., Herz, D. M., Brown, P., Forstmann, B. U. & Zaghloul, K. Frontosubthalamic circuits for control of action and cognition. in Journal of Neuroscience 36, 11489–11495 (2016).

43. Forstmann, B. U. et al. Cortico-subthalamic white matter tract strength predicts interindividual efficacy in stopping a motor response. Neuroimage 60, 370–375 (2012).

44. Deco, G., Jirsa, V. K. & McIntosh, A. R. Emerging concepts for the dynamical organization of resting-state activity in the brain. Nature Reviews Neuroscience 12, 43–56 (2011).

45. Greicius, M. D., Supekar, K., Menon, V. & Dougherty, R. F. Resting-State Functional Connectivity Reflects Structural Connectivity in the Default Mode Network. Cereb. Cortex 19, 72–78 (2009).

46. Nambu, A. Globus pallidus internal segment. Progress in Brain Research 160, 135–150 (2007).

47. Quartarone, A. et al. New insights into cortico-basal-cerebellar connectome: clinical and physiological considerations. Brain 143, 396–406 (2020).

48. Cassidy, M. & Brown, P. Spectral phase estimates in the setting of multidirectional coupling. J. Neurosci. Methods 127, 95–103 (2003).

49. Swann, N. C. et al. Elevated synchrony in Parkinson disease detected with electroencephalography. Ann. Neurol. 78, 742–750 (2015).

50. Jackson, N., Cole, S. R., Voytek, B. & Swann, N. C. Characteristics of waveform shape in Parkinson’s disease detected with scalp electroencephalography. eNeuro 6, (2019).

51. Sherman, M. A. et al. Neural mechanisms of transient neocortical beta rhythms: Converging evidence from humans, computational modeling, monkeys, and mice. Proc. Natl. Acad. Sci. U. S. A. 113, E4885–E4894 (2016).

52. Burke, J. F., Ramayya, A. G. & Kahana, M. J. Human intracranial high-frequency activity during memory processing: Neural oscillations or stochastic volatility? Current Opinion in Neurobiology 31, 104–110 (2015).

53. Lozano-Soldevilla, D., Huurne, N. & Oostenveld, R. Neuronal oscillations with non-sinusoidal morphology produce spurious phase-to-amplitude coupling and directionality. Front. Comput. Neurosci. 10, (2016).

54. Fogelson, N. et al. Different functional loops between cerebral cortex and the subthalmic area in Parkinson’s disease. Cereb. cortex 16, 64–75 (2006).

55. Li, Q. et al. Therapeutic deep brain stimulation in Parkinsonian rats directly influences motor cortex. Neuron 76, 1030–41 (2012).

56. Malekmohammadi, M. et al. Pallidal stimulation in Parkinson disease differentially modulates local and network β activity. J. Neural Eng. 15, 056016 (2018).

57. Silberstein, P. et al. Patterning of globus pallidus local field potentials differs between Parkinson’s disease and dystonia. Brain 126, 2597–608 (2003).

58. Neumann, W. J. et al. Subthalamic synchronized oscillatory activity correlates with motor impairment in patients with Parkinson’s disease. Mov. Disord. 31, 1748–1751 (2016).

59. Foley, J. A., Foltynie, T., Limousin, P. & Cipolotti, L. Standardised Neuropsychological Assessment for the Selection of Patients Undergoing DBS for Parkinson’s Disease. Parkinsons. Dis. 2018, (2018).

60. Hartmann, C. J., Fliegen, S., Groiss, S. J., Wojtecki, L. & Schnitzler, A. An update on best practice of deep brain stimulation in Parkinson’s disease. Ther. Adv. Neurol. Disord. 12, 175628641983809 (2019).

61. Ramirez-Zamora, A. & Ostrem, J. L. Globus pallidus interna or subthalamic nucleus deep brain stimulation for Parkinson disease a review. JAMA Neurology 75, 367–372 (2018).

62. Lalo, E. et al. Patterns of bidirectional communication between cortex and basal ganglia during movement in patients with Parkinson disease. J. Neurosci. 28, 3008–16 (2008).

63. Horn, A. et al. Connectivity Predicts deep brain stimulation outcome in Parkinson disease. Ann. Neurol. 82, 67–78 (2017).

64. Fox, M. D. Mapping symptoms to brain networks with the human connectome. N. Engl. J. Med. 379, 2237–2245 (2018).

65. Gibb, W. R. & Lees, A. J. A comparison of clinical and pathological features of young- and old-onset Parkinson’s disease. Neurology 38, 1402–6 (1988).

66. Gratwicke, J. et al. Bilateral deep brain stimulation of the nucleus basalis of meynert for Parkinson disease dementia a randomized clinical trial. JAMA Neurol. 75, 169–178 (2018).

67. Horn, A., Ostwald, D., Reisert, M. & Blankenburg, F. The structural-functional connectome and the default mode network of the human brain. Neuroimage 102, 142–151 (2014).

68. Mitra, P. P. & Pesaran, B. Analysis of dynamic brain imaging data. Biophys. J. 76, 691–708 (1999).

69. Gross, J. et al. Dynamic imaging of coherent sources: Studying neural interactions in the human brain. Proc. Natl. Acad. Sci. U. S. A. 98, 694–9 (2001).

