## Supplementary Materials for "Neural signatures of pathological hyperdirect pathway activity in Parkinson’s disease"

**Supplementary Methods**

**Surgical Procedure**

A Medtronic (Medtronic Neurological Division, Minneapolis, MN) model 3389 electrode was implanted in all but one subject (GPi 1), who was implanted with the Medtronic model 3387 electrode. Each electrode comprised four platinum-iridium contacts (model 3389 – diameter 1.27 mm, length 1.5 mm, centre-centre separation 2 mm; model 3387 – diameter 1.27 mm, length 1.5 mm, centre-centre separation 3 mm) that are numbered from 0 (lowermost) to 3 (uppermost). The surgical targets investigated here were the dorsal motor region of the STN and the posterior third of the ventral pallidum in the two cohorts. Further details of the operative procedure can be found in ^1,2^.

The locations of the electrodes were confirmed following implantation with immediate postoperative fast spin-echo T2-weighted magnetic resonance imaging (MRI) with a Leksell frame still in situ. Stainless steel electrode extension cables were externalized through the scalp to enable recordings prior to connection to a subcutaneous DBS pacemaker, implanted in a second operative procedure seven days later. Recordings were performed 3–6 days after electrode implantation. Study procedures were approved by the National Research Ethics Service Committee South Central – Oxford B, and the patients gave written informed consent prior to participation.

**Simultaneous magnetoencephalography and local field potential recordings**

MEG recordings were performed using a CTF 275-channel MEG system (CTF/VSM MedTech). MEG data were sampled at 2400 Hz and stored to disk for subsequent offline analyses. LFP activity recorded from DBS electrodes was collected at the same time as MEG using a battery-powered and mains optically isolated BrainAmp system (Brain Products). Three bipolar channels (0-1, 1-2, 2-3) were recorded from each electrode and were high-pass filtered at 1 Hz in the hardware to avoid amplifier saturation due to large DC offsets.

Recordings were performed at rest whilst patients were on their usual medication. The patient was requested to stay still during recordings and to keep their eyes open whilst focussing on a fixation cross that appeared on a screen in front of them. Rest recordings had a duration of 3 minutes. A neurologist was present in the magnetically shielded room at all times in order to ensure patient safety.

**Generation of volumetric coherence images using beamforming**

Beamforming relies on a linear projection of sensor data using a spatial filter that is computed from the lead-field of a location of interest and either the data covariance or the cross-spectral density matrix ^3,4^. Lead-fields were computed using a single shell head model ^5^. The model was generated in SPM12 based on the patient’s preoperative structural MRI and fiducial-based co-registration was performed.

The source space was defined as a 5 mm spaced grid in MNI space bounded by the inner skull surface. The resulting values of coherence – between each grid point and the subcortical LFP - were linearly interpolated to produce 3D volumetric images with 2 mm resolution for visualisation. Coherence images were smoothed with an 8 mm isotropic Gaussian kernel to ensure conformance to the assumptions of random field theory prior to statistical analysis in SPM.

**Estimation of directionality**

The effective directionality of coupling between the cortex and the STN/GPi LFP was computed with a non-parametric variant of spectral Granger causality ^6,7^. To determine the significance of directionality estimates, we compared the Granger estimate of original data to that of surrogate time-reversed data using a paired *t*-test ^8^. Taking the example of two signals A and B, with A Granger causing B, the Granger causality from A to B should be higher for the original than for the time-reversed data, giving rise to a positive difference. In contrast, the estimate of causality from B to A should be increased by time reversal thereby giving a negative difference. In the event of there being statistically significant coupling in one particular direction (e.g. cortex to sub-cortex), time delays were computed by regressing the unwrapped angle of the cross-spectrum between the two component signals against frequency ^9^. Time delays were statistically compared for the STN and GPi with covariates to account for both subject-specific dependencies in the recordings from the two hemispheres (between subjects), and for potential differences between the recordings from the right and left sides in individual subjects (within subjects). Covariates representing each patient’s preoperative levodopa equivalent dose, UPDRS Part III motor score on medication and MMSE were also introduced in order to account for phenotypic differences between the two groups.

**Waveform clues to distinct origins**

The sharpness of neural population activity is increasingly thought to be associated with the tight synchronisation of a regular input or drive, so that the temporal summation of synaptically driven polarisation changes gives sharp transients ^10–15^. Thus far this has only been proposed at the cortical level and on theoretical rather than empirical grounds. Here we investigated if the sharpness of waveform extremata in the STN or GPi distinguished between the two nuclei and supported a more synchronised input to the former.

As above, only contact pairs where at least one contact lay in the target nucleus were considered. Off-line LFP activity was down-sampled to 1kHz, notch filtered at 50 Hz and its harmonics, and high-pass ﬁltered at 3 Hz. To improve the signal-to-noise ratio of the beta component in the LFP we extracted bursts of higher power using methods that have been previously outlined ^16^. Specifically, LFP signals were decomposed using Wavelet transformation using the FieldTrip toolbox (<http://www.fieldtriptoolbox.org/>; fieldtrip-function ft_freqanalysis, morlet wavelet width = 10, gwidth = 5) into frequencies ranging from 1 to 40 Hz with a resolution of 1 Hz. The frequency of the maximum amplitude bin in the beta frequency range (1-Hz bins between 13 and 30 Hz) was selected and the corresponding time evolved wavelet amplitude (bandwidth = 5 Hz) was smoothed (0.2 sec) and DC corrected (by subtracting the non-overlapping 20-sec moving average). Time points at which the time evolved wavelet amplitude exceeded a fixed amplitude threshold deﬁned as the 75th percentile amplitude were determined as samples within beta bursts. Next we selected for further analysis bursts with a duration of 400ms.

To investigate the shape of the waveforms comprising beta bursts in the STN or GPi, we used empirical mode decomposition (EMD) to decompose a signal into elementary signals referred to as intrinsic mode functions (IMFs) ^17^. To calculate sharpness within beta bursts, IMFs with peak power in the beta frequency band were extracted using ensemble EMD (level of added noise = 0.1, ensemble number = 100) with the influences from the added noise less than a fraction of 1% of the standard deviation. The IMF with the highest peak power within the beta band was selected as the beta IMF. The sharpness of each extrema in the beta IMF was defined as the average of the two absolute voltage differences between the extrema and the 5 ms either side of that point ^13^. The sharpness of all extrema within each beta burst were averaged, and then these values averaged again to give a single sharpness value for each LFP. Note that we considered the sharpness of extrema (peaks and troughs) independently of their polarity, as the latter is determined by the orientation of the contact pair used to record the LFP with respect to the generator. Accordingly, sign may flip between recordings in an unconstrained way, so here we averaged the sharpness of trough and peaks. As sharpness may depend on signal-to-noise ratios and frequencies of oscillations we also estimated sharpness after STN and GPi LFP signals were matched for power and the centre frequencies of beta-band peaks were normalised to a fixed period of 50 ms.

**Computational modelling of high and low beta band oscillatory synchrony**

The model developed here rests on the premise that the change in the average firing rate of a population *i*,
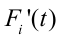
is governed by the population time constant,
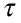
 the current firing rate,
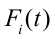
 and a sigmoidal function,
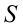
 of the sum of all delayed synaptic inputs,
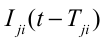
from sources *j* to population *i* multiplied by their respective connection strengths,
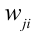
 ^18^:

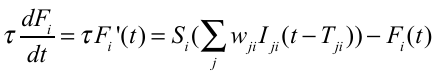

The sigmoid function, *S_i_* in (1) represents the activation function of population *i* in response to synaptic inputs and is parameterised by the maximal firing rate of the population (
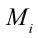
) and its firing rate in the absence of inputs (
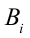
) as indicated for each population in (3).

Additional features which define the sigmoid function are the firing rate in response to zero input and the maximal slope. We modelled the dynamics of five neuronal populations within the cortico-basal-ganglia circuit: 1) An excitatory cortical population, E ,2) an inhibitory cortical population, I, 3) the STN, 4) the GPe and 5) the GPi. **Supplementary Figure 6A** provides an illustration of modelled populations and their connections. In keeping with the work of Pavlides et al., 2015 cortical activity was modelled with an excitatory and inhibitory population, with the former providing inputs to the basal ganglia. Additionally in the present model we included: 1) an auto-inhibitory self-connection in the inhibitory cortical neural population which has been considered in previous accounts of cortical Jansen-Rit type neural mass models ^19^, 2) an explicit model of the dynamics of the GPi based on its anatomical connectivity ^20,21^ as our primary motive was to compare the activity and connectivity profile of the STN and GPi and 3) an additive signal dependent stochastic noise term so that the differential equations describing the evolution of firing rates for each population were stochastic delay differential equations (SDDEs) ^22,23^. The addition of signal dependent noise serves to ensure that the spike rate variance correlates with the mean firing rate at each integration step,
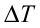
 (see for example Dayan and Abbott, 2001 for a discussion of the Fano factor).

Rather than explicitly modelling the firing dynamics of the striatum we considered inhibitory striatal inputs, *S* to the GPi and GPe to be fixed in keeping with ^24^. Similarly the excitatory cortical population, *E* receives a constant component of intrinsic and extrinsic excitatory inputs, *C*. We also consider that the net effect of excitation of the GPi-thalamo-cortical loop has an inhibitory effect on excitatory cortical population activity. The equations governing the dynamics of the cortico-basal-ganglia circuit in our model are as follows, where
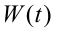
represents the standard Weiner process:

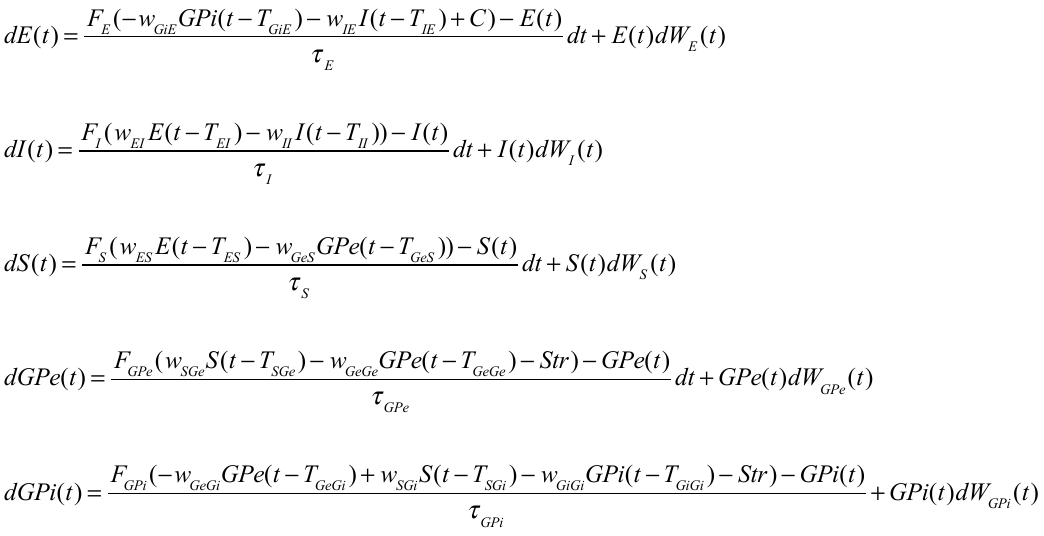
The sigmoid functions governing the activation function of each population are as follows:

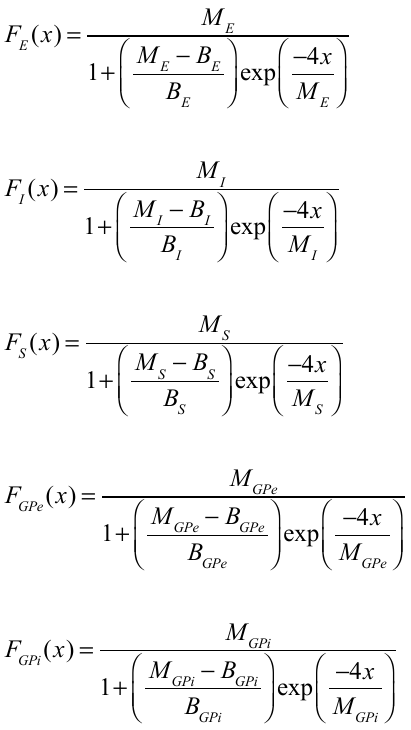

The resulting SDDEs were integrated over a time period of 4 seconds using a Euler-Maruyama integration scheme ^22^ with a step size,
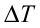
 of 10^-4^ seconds where the additive noise term is given by:
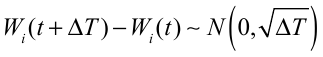
.

Simulations with noise were performed to ensure physiological values of coherence (**Figure 7**). However, we also investigate the behaviour of this system without noise by integrating the above equations in the absence of the noise term using the built-in MATLAB integrator, DDE23 (**Figure 6 and** **Supplementary Figure 6**). In contrast to ^24^ who focus purely on the generation of broad band beta oscillations our aim was to capture mechanisms underlying the generation of low and high beta band activities that could explain both our data and those of other studies. Where possible, parameters of our model were selected from those previously detailed in ^24^ (see **Supplementary Table 2** for further details of parameters used).

**Supplementary Results**

**Supplementary analysis of cortico-STN and cortico-GPi delays in the high beta frequency band**

In the main results we show that time delays between cortex and the STN in the high beta frequency band are shorter than those between cortex and the GPi across the entire beta frequency range. We motivated using different bands for the two subcortical structures based on spectral differences in their coherence profiles with the cortex. Importantly however, the longer time delay to GPi persisted if the above analysis was repeated whilst limiting both coupling to STN and GPi to the upper beta frequency band (see **Supplementary Figure 3**; in this case there was also only a significant main effect of target location, F(1,32) =10.1, P <0.01).

**Non-sinusoidality of the beta waveform supports**

It was reasoned that the sharpness of LFP extrema, thought to reflect tight synchronisation of inputs, might also provide orthogonal support for dominance of a very direct pathway from cortex to STN, and a further distinction between STN and GPi LFPs ^10–15^. In the case of the STN LFP tight synchronisation might reflect precise timing at the cortical level and/or lack of temporal dispersion in monosynaptic hyperdirect pathways. An analysis of LFP sharpness confirmed that the extrema of LFP beta oscillations were sharper in the STN than in the GPi (see bars labelled ‘unmatched’ in **Supplementary** **Figure 4A**; *F*(1,60) =16.2, *P < 1x*10^-3^). This difference was maintained if sharpness was estimated after LFP signals were matched for power and after the centre frequencies of beta-band peaks were normalised to a fixed period (see bars labelled ‘matched’ in **Supplementary** **Figure 4A**; *F*(1,26)= 4.9, *P* < 0.036).

Finally we reasoned that if STN waveform sharpness is reflective of tight synchronisation with cortical sources in the monosynaptic hyperdirect pathway, one may also expect to observe a relationship between hyperdirect tract density and beta band extrema sharpness. In keeping with this hypothesis, for the STN our results reveal a significant correlation between LFP sharpness and hyperdirect fibre tract density (**Supplementary Figure 4B**; r^2^ correlation coefficient = 0.61, p = 0.02). This was not the case for the GPi (r^2^ = 0.09, p = 0.5).

**Simultaneous visualisation of cortico-subcortical fibre tracts and MEG networks**

**Supplementary Figure 5** displays the intersection of tractography derived structural connectivity and MEG derived functional connectivity. The magenta streamlines indicate tracts passing between the broadband beta network (shown in yellow) and the STN (left panel) and GPi (right panel). The cyan streamlines indicate tracts passing between the high beta network (shown in red) and the STN and GPi. Since voxels comprising the high beta network were contained within those comprising the broadband beta network, the former were subtracted from the latter to highlight the difference in associated tracts. The colour bars indicate the number of repetitions of each fibre across subjects.

**Computational model: cortically generated high beta frequency activity can induce the generation of lower frequency beta rhythms in the subcortex.**

Our model was set up to account for the observation that high beta activity generated in the cortex propagates subcortically to the STN (see left image in **Supplementary Figure 6A** for the structure of the model). This is in keeping both with our present analysis of cortico-STN directionality in the high beta frequency range and with the results of other reports ^9,25,26^. The middle and right most images in **Supplementary** **Figure 6A** display the cortical generation of high beta frequency oscillations. In this simulation we consider only the cortical excitatory (E) and inhibitory (I) populations and therefore the top-down and bottom-up connections (W_ES_ and W_GiE_) to and from the basal ganglia are set to zero. The middle image in (A) details the effect of changing the coupling parameters, W_IE/EI_ on the peak frequency generated by the cortical populations. The right most image in (A) shows simulated firing rates and corresponding spectra for the cortical populations with the coupling parameter, W_IE/EI_ set to a value of 4. In this instance both the E and I populations produce oscillatory activity with a frequency of approximately 27 Hz. The remaining parameters in this simulation are as per **Supplementary Table 2**.

Secondly, in our model low beta frequencies are generated by delayed interactions in the reciprocal loop between the STN and the GPe. It has previously been shown that this loop is capable of producing oscillations under certain conditions when the system transitions from a stable to an unstable fixed point through a Hopf bifurcation ^24,27^. **Supplementary** **Figure 6B** shows a simulation similar to that performed in **Supplementary** **Figure 6A**, but this time with the introduction of top-down (W_ES_) and bottom-up connections (W_GiE_) to and from the basal ganglia. We use fixed values of W_ES_ and W_EI/IE_ but vary the strength of the net inhibitory loop between the GPi, thalamus and cortex, W_GiE_. In each of the three images in **Supplementary** **Figure 6B**, the top subplot displays the integrated time series, whilst the middle subplot shows the resulting power spectra and the bottom subplot a phase portrait of STN and GPe firing rates after the system reaches the vicinity of a periodic orbit. In these phase portraits activity of GPe within a low beta oscillation cycle is plotted against the activity in STN, and different points on the curve correspond to different time points within a cycle. The phase portraits are colour coded depending on the input excitatory cortical population firing rate, thus the two yellow segments of the curve correspond to peaks of cortical high beta oscillation. At low values of W_GiE_ the time series of STN, GPi and GPe comprise high and low beta frequency components (best seen in the left image in **Supplementary** **Figure 6B**) that are observed in the time series, the spectra and the profile of the STN/GPe phase portraits. In summary, the model including two generators of oscillation in cortex and STN-GPe circuit qualitatively reproduced the two peaks in beta activity seen in **Figures 2 and 3**.

**Comment on bursting behaviours**

Recent work has focussed on the importance of transient rather than sustained episodes of synchrony being important in both pathological and physiological states ^16,28,29^. Although the model presented here does not explicitly focus on the generation of bursting behaviours, it is easy to see how bursts may arise. Endogenous fluctuations (triggered by noise) of the inputs to the reciprocal STN-GPe loop when it is operating close to its bifurcation point may trigger transient oscillatory behaviours (see the left image in **Figure 6B**). This model therefore makes the testable prediction that transient bursts of cortical high beta activity can trigger the generation of lower beta frequency bursts within the STN. The genesis of bursting activity within similar models is likely to be a focus of future work.

**Supplementary Figures**

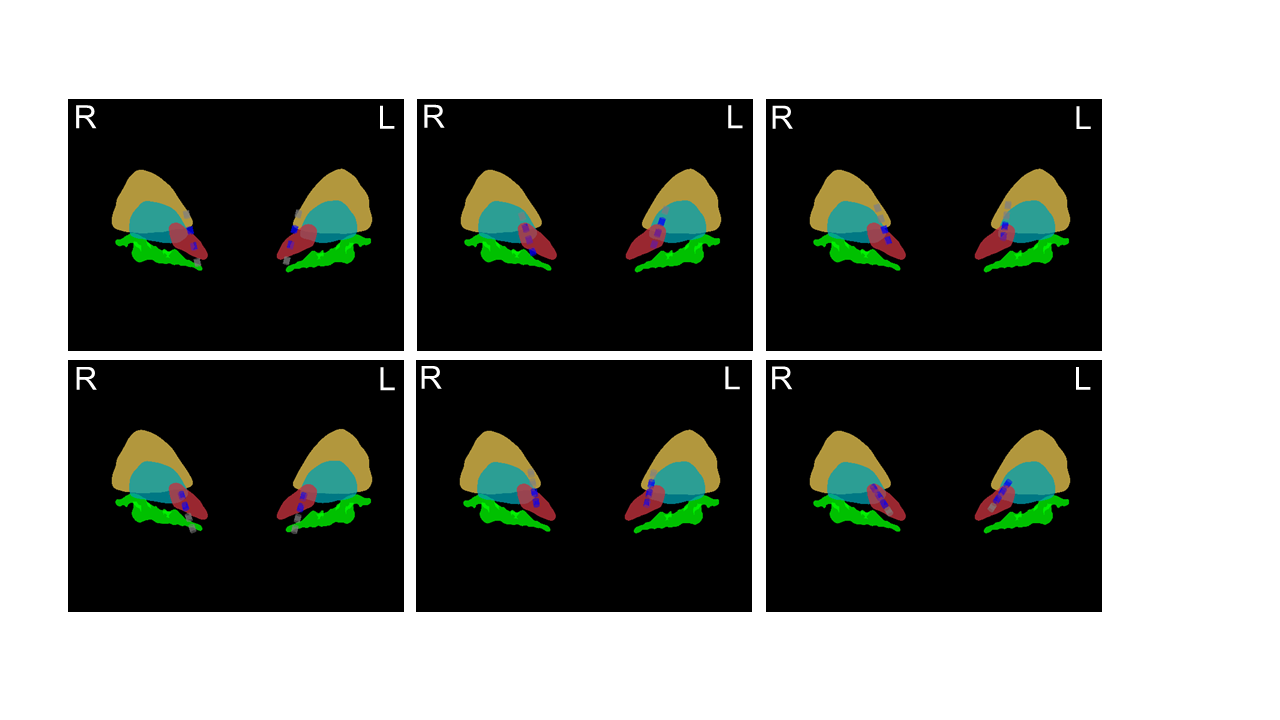

**Supplementary Fig 1. Individual STN contact localisations**

All STN contacts are visualised in MNI template space in the coronal plane, individually for each of the six STN DBS patients studied in this report. Templates of the STN (red), GPi (turquoise), external segment of the Globus Pallidum (GPe; yellow) and nucleus basalis of Meynert (NBM; green) are shown. Contacts located partially or completely within the STN template are coloured in blue, whilst those located outside the STN are coloured grey.

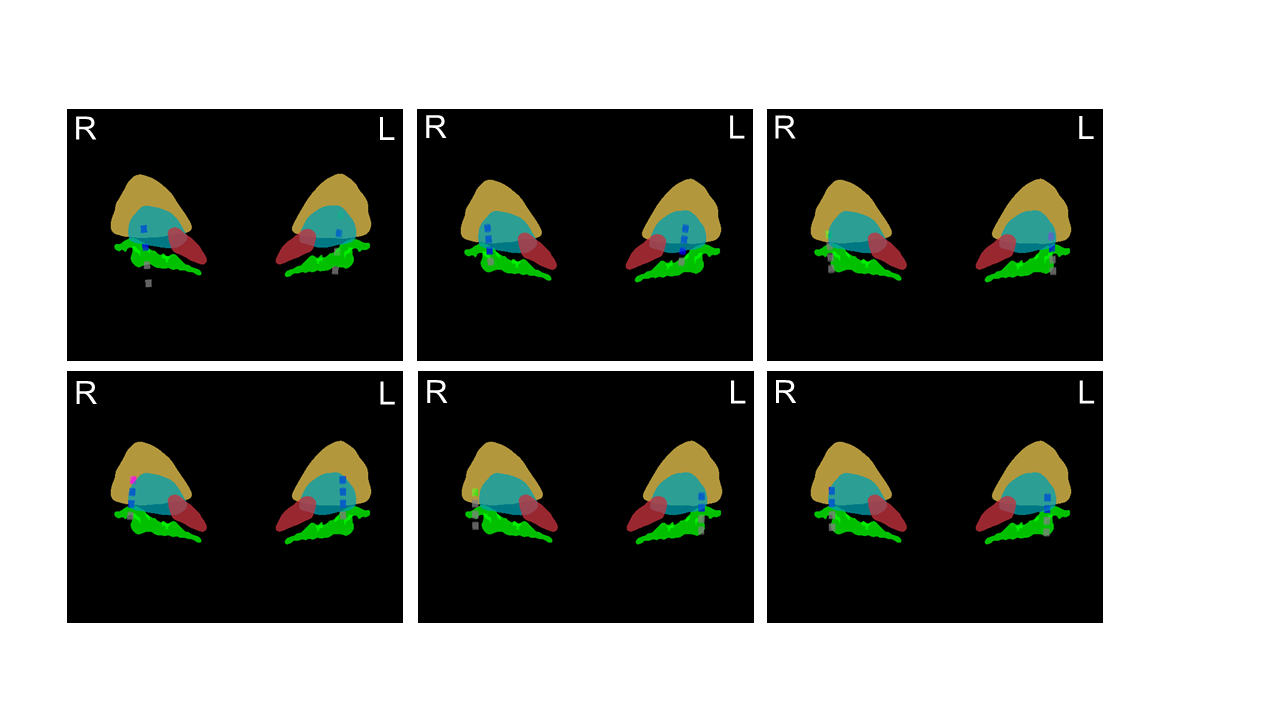

**Supplementary Fig 2. Individual GPi contact localisations**

All GPi contacts are visualised in MNI template space in the coronal plane, individually for each of the six GPi DBS patients studied in this report. Templates of the STN (red), GPi (turquoise), external segment of the Globus Pallidum (GPe; yellow) and nucleus basalis of Meynert (NBM; green) are shown. Contacts located partially or completely within the GPi template are coloured in blue, whilst those located in the GPe are coloured green. One contact traversed both structures and is coloured magenta.

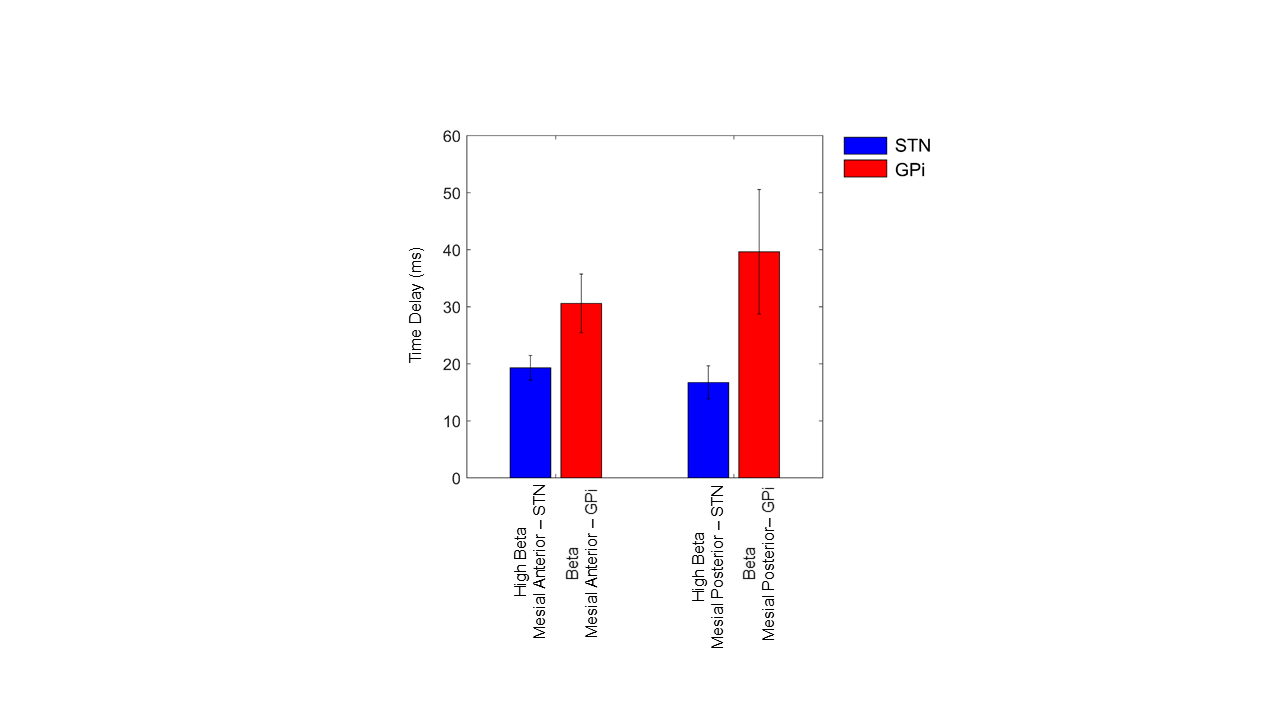

**Supplementary Fig 3. Estimation of time delays between cortex and the STN/GPi**

Time delays between cortex and the STN/GPi were estimated across the high beta frequency range (21-30 Hz). Vertical bars represent standard errors of the mean. See Supplementary Results for further discussion.

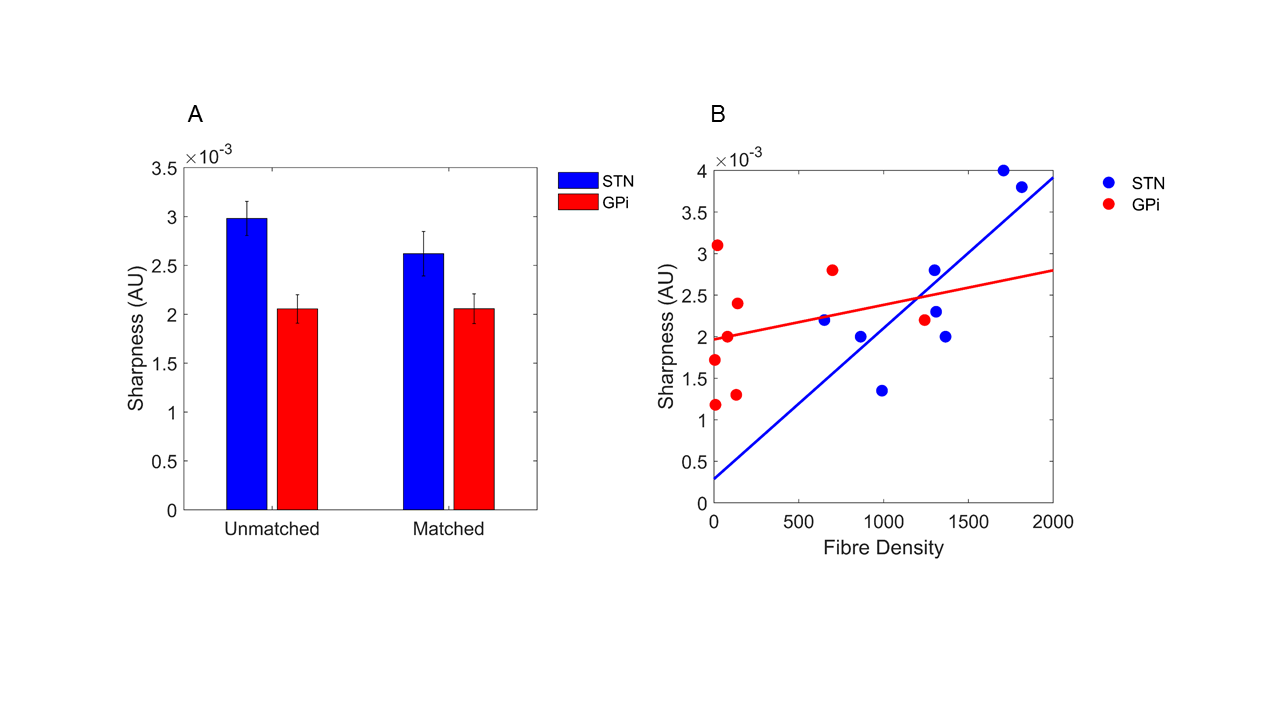

**Supplementary Fig 4. Relationship between waveform shape and structure: sharpness is greater in STN and predicted by tract density.** (A) The sharpness of the extrema of LFP beta oscillations is greater in the STN than in the GPi (*F*(1,60) =16.2, *P < 1x*10^-3^). Data from all contacts in each target are considered (see supplementary figures 1 and 2). Vertical bars represent standard errors of the mean. The same is true even if beta IMFs are matched for power and frequency (*F*(1,26)= 4.9, *P* < 0.036). (B) Fibre tract density in the hyperdirect pathway is significantly predictive of waveform sharpness within the STN (r^2^ = 0.61, p = 0.02). In contrast cortico-GPi tract densities did not correlate with GPi waveform sharpness (r^2^ = 0.09, p = 0.5). The straight lines are linear regression fits. Results in (B) are matched for both frequency and power. After exclusion of STN or GPi contact pairs due to inability to match for power, data from all target contact pairs within each hemisphere were averaged to yield a single value for each hemisphere for visualisation. Data from a total of 8 STN and 8 GPi hemispheres is shown (data from 4 STN and 4 GPi hemispheres were rejected due to inability to match for power). Results were similar if the correlation was performed for all contact pairs without matching for power or frequency (data not shown R^2^ = 0.67, p = 0.02 in STN; R^2^ = 0.23, p = 0.20 in GPi).

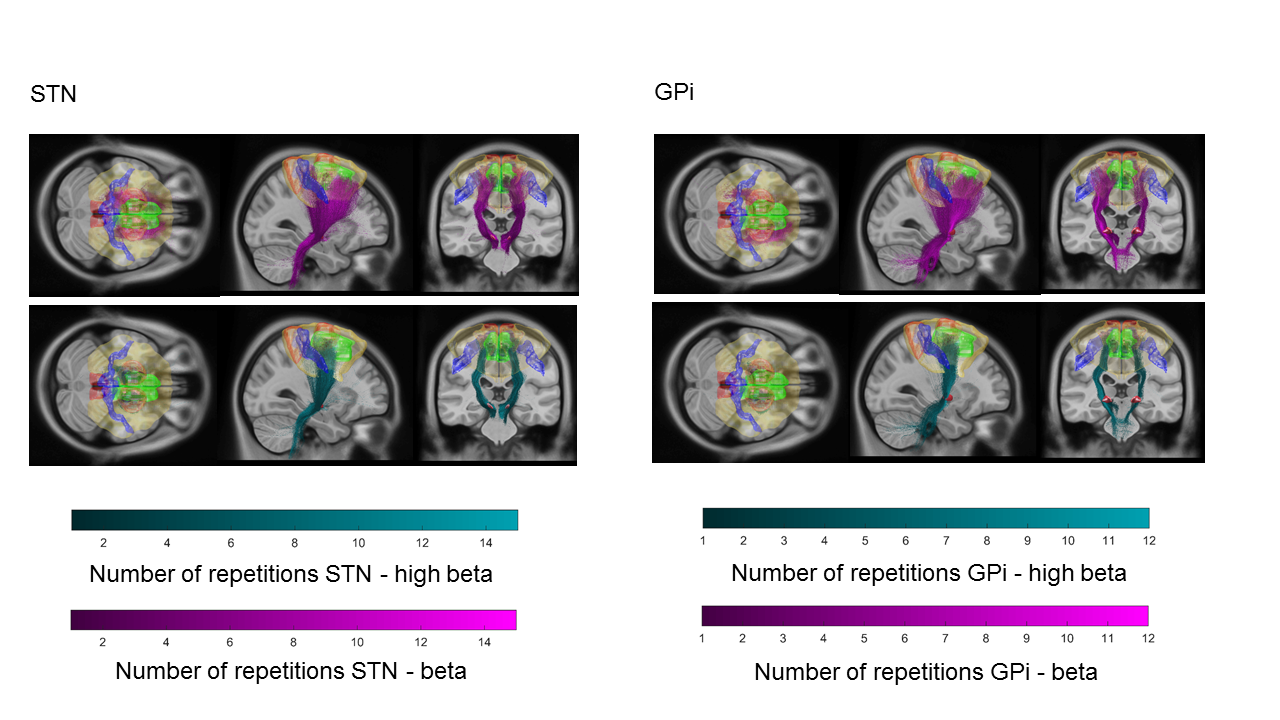

**Supplementary Fig 5. Intersection of group level MEG derived cortico-STN/cortico-GPi networks and tractography streamlines**. Cortico-STN and cortico-GPi networks derived from concurrent MEG and LFP recordings are displayed together with fibre streamlines on a T1-weighted MRI scan. The left panel depicts fibres passing to STN contacts, whilst the right panel shows fibres passing to GPi contacts. In each case, fibres were selected to originate in either: 1) cortical regions where there was a significant main effect of band for the STN (termed the high beta network which is shown in red with fibres originating from it coloured cyan) or 2) cortical regions that couple to the STN and GPi across the entire beta frequency range (termed the beta network which is shown in yellow with fibres originating from it coloured in magenta). Fibres are colour coded depending on the number of times they were repeatedly chosen (see colour bar). Cortical regions shown in blue and green, indicate boundaries of the primary motor cortex and SMA respectively.

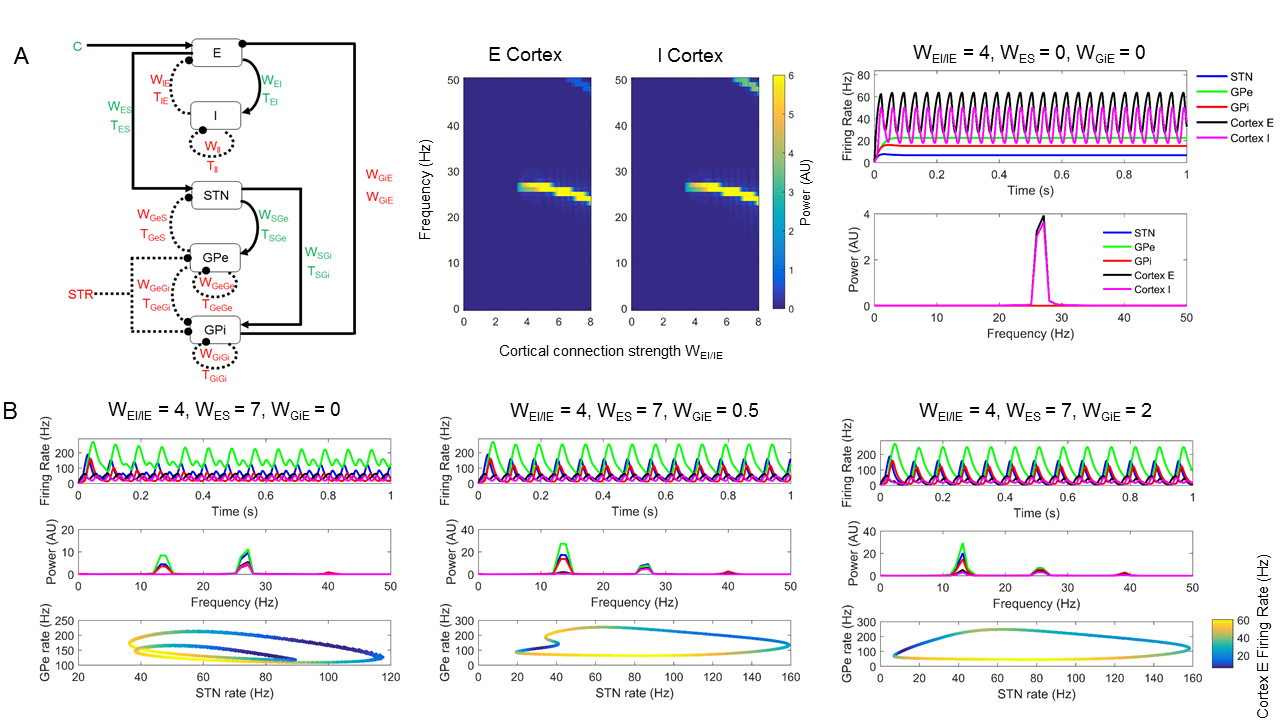

**Supplementary Fig 6. Computational model of the generation of cortical high beta frequency activity which can in turn trigger resonance at lower beta frequencies within the STN-GPe loop.** (A): the left image displays the connectivity of the cortico-basal-ganglia circuit in the computational model. The computational model developed includes the major connections of the basal ganglia. Excitatory connections are depicted by filled black lines with arrows, whilst inhibitory connections are indicated by dashed black lines with circles. E = cortical excitatory population, I = cortical inhibitory population, S = STN, Ge = GPe and Gi = GPi. W_ES_ and T_ES_ indicate the strength and delay of synaptic connections between the excitatory cortical population and the STN. Inhibitory inputs, connections and delays are coloured red, whilst excitatory inputs, delays and connections are coloured green. Inhibitory striatal (Str) inputs to GPe and GPi are considered to be fixed in our model. An input, C representing a constant component of intrinsic and extrinsic excitatory inputs is provided to the excitatory cortical population, E. The inhibitory cortical population, I, the GPe and the GPi have self-inhibitory connections with delays. Differential equations governing the dynamics of studied nodes are detailed in equation 1. The middle and right images of (A) reveal the behaviours of the excitatory cortical population (E), the inhibitory cortical population (I), the STN, the GPe and the GPi in the absence of top-down and bottom-up connections to and from the basal ganglia. This is equivalent to setting, W_ES_ and W_GiE_ equal to 0. Simulations are performed in the absence of additive noise. The middle image in (A) shows the effect of changing the coupling parameters, W_IE/EI_ on the peak frequency generated by the cortical populations. The right image in (A) shows simulated firing rates and corresponding spectra for the cortical populations with the coupling parameter, W_IE/EI_ set to a value of 4. In this simulation, the remaining parameters were set to those detailed in **Supplementary Table 2**. With W_EI_ and W_IE_ set to 4 the cortical population is capable of generating oscillatory activity in the high beta frequency range (approximately 27 Hz). In Panel (B) top-down (W_ES_) and bottom-up connections (W_GiE_) to and from the basal ganglia are introduced. We use fixed values of W_ES_ and W_EI/IE_ but vary the strength of the net inhibitory loop between the GPi, thalamus and cortex (W_GiE_). For each of the three images in panel B, the top subplot shows the integrated time series (firing rates) for the excitatory cortical population (E), the inhibitory cortical population (I), the STN, the GPe and the GPi. The middle subplot shows power spectra of the integrated time series and the bottom subplot shows a phase portrait of the STN and GPe activities colour coded by the firing rate of the excitatory cortical population, E, which provides inputs to the STN. At low values of W_GiE_ the time series of STN, GPi and GPe comprise high and low beta frequency components which are also reflected in the periodic orbits of the phase portraits.

**Supplementary Tables**

| **Case** | **Age** | **Parkinson’s**  **Disease Duration**  **(Years)** | **Preoperative Medication (mg)** | **Unified Parkinson’s**  **Disease Rating scale - UPDRS III**  **On medication** | **Cognitive Score**  **(MMSE)** |
| --- | --- | --- | --- | --- | --- |
| STN 1 | 51 | 8 | LDE 650 | 21 | 29 |
| STN 2 | 54 | 8 | LDE 2150 | 9 | 29 |
| STN 3 | 66 | 6 | LDE 404 | 18 | 30 |
| STN 4 | 58 | 11 | LDE 1320 | 25 | 30 |
| STN 5 | 54 | 15 | LDE 1150 | 19 | 29 |
| STN 6 | 60 | 17 | LDE 1220 | 13 | 29 |
| **Mean STN** | **57.2** | **10.8** | **1149** | **17.5** | **29.3** |
| GPi 1 | 61 | 14 | LDE 500 | 27 | 25 |
| GPi 2 | 75 | 11 | LDE 670 | 26 | 24 |
| GPi 3 | 73 | 15 | LDE 380 | 42 | 25 |
| GPi 4 | 65 | 11 | LDE 923 | 33 | 25 |
| GPi 5 | 46 | 10 | LDE 575 | 37 | 22 |
| GPi 6 | 71 | 15 | LDE 833 | 16 | 21 |
| **Mean GPi** | **65.2** | **12.7** | **647** | **30.2** | **23.7** |

**Supplementary Table 1. Clinical characteristics of patients**. LDE - levodopa dose equivalent. The total UPDRS III ^30^motor score is presented on medication. Cognitive function was assessed with the the Mini-Mental State Examination, MMSE ^31^.

| **Parameter** | **Value** | **Comment/Reference** |
| --- | --- | --- |
| **Fixed inputs** | |  |
| C | 172.18 | ^24^ |
| Str | 8.46 | ^24^ |
| **Synaptic connection weight (AU)** | |  |
| W_IE_/W_IE_ | 4 | ^24^ |
| W_ES_ | 7 | ^24^ |
| W_II_ | 3 | We reasoned that strength of auto-inhibition might be less than that of inhibition of excitatory population as per ^19^ who study cortical columns in a Jansen-Rit model |
| W_GeS_ | 1.3 | ^24^ |
| W_SGe_ | 4.87 | ^24^ |
| W_GeGe_ | 0.53 | ^24^ |
| W_SGi_ | 4 | We expect that this would be similar to W_SGe_ based on tractography and cellular studies of STN efferents to GPi/GPe ^32–34^ |
| W_GeGi_ | 1 | We reasoned that this would be stronger than W_GeGe_ based on higher density of synapses from GPe neurones to GPi neurones than to GPe neurones ^33^ |
| W_GiGi_ | 0.53 | We account for the possibility that GPi neurones may inhibit their own activation via recurrent connections ^33,35^ |
| W_GiE_ | 0.5 | We explored a number of values for this parameter and conclude that the net inhibitory effect of activation of the GPi-thalamo-cortical loop is small in keeping with previous studies ^36–38^ |
| **Delays (ms)** | |  |
| T_IE_/T_IE_ | 5 | Mean of range previously reported ^24^ |
| T_II_ | 4 | We reasoned that auto-inhibitory delays between cells of the same population are fixed. We therefore used constant values for T_II_, T_GiGi_ &T_GeGe_ |
| T_ES_ | 5.5 | ^24^ |
| T_GeS_ | 6 | ^24^ |
| T_SGe_ | 6 | ^24^ |
| T_GeGe_ | 4 | ^24^ |
| T_SGi_ | 6 | Similar to T_GeS_ based on evoked responses to STN stimulation ^39^ |
| T_GeGi_ | 6 | Similar to T_SGe_ based on latency of inhibitory response of STN→GPe→GPi following single pulse stimulation of STN ^39^ |
| T_GiGi_ | 4 | We reasoned that auto-inhibitory delays between cells of the same population are fixed. We therefore used constant values for T_II_, T_GiGi_ &T_GeGe_ |
| T_GiE_ | 20 | Within the range delays previously reported  ^40^ |
| **Time constants (ms)** | |  |
| 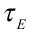 | 11 | ^24^ |
| 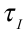 | 11 | ^24^ |
| 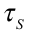 | 12.8 | ^24^ |
| 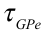 | 20 | ^24^ |
| 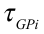 | 14 | ^21^ |
| **Sigmoid function firing parameters (spikes/s)** | |  |
| M_E_/B_E_ | 75/17.85 | ^24^ |
| M_I_/B_I_ | 205/9.87 | ^24^ |
| M_s_/B_s_ | 300/10 | ^24^ |
| M_GPe_/B_GPe_ | 400/10 | ^24^ |
| M_GPi_/B_GPi_ | 400/18 | Taken from in vitro recordings and responses to stimulation ^21,39^ |

**Supplementary Table 2. Parameter values used for generating simulated data from the computational model.** The majority of parameter values are derived from ^24^. Where we have introduced additional parameters, we provide a comment or reference justifying selection of the particular value.

**References**

1.	Foltynie, T. & Hariz, M. I. Surgical management of Parkinson’s disease. *Expert Rev. Neurother.* **10,** 903–14 (2010).

2.	Foltynie, T. *et al.* MRI-guided STN DBS in Parkinson’s disease without microelectrode recording: efficacy and safety. *J. Neurol. Neurosurg. Psychiatry* **82,** 358–63 (2011).

3.	Van Veen, B. D., van Drongelen, W., Yuchtman, M. & Suzuki, A. Localization of brain electrical activity via linearly constrained minimum variance spatial filtering. *IEEE Trans. Biomed. Eng.* **44,** 867–80 (1997).

4.	Gross, J. *et al.* Dynamic imaging of coherent sources: Studying neural interactions in the human brain. *Proc. Natl. Acad. Sci. U. S. A.* **98,** 694–9 (2001).

5.	Nolte, G. The magnetic lead field theorem in the quasi-static approximation and its use for magnetoencephalography forward calculation in realistic volume conductors. *Phys. Med. Biol.* **48,** 3637–52 (2003).

6.	Brovelli, A. *et al.* Beta oscillations in a large-scale sensorimotor cortical network: directional influences revealed by Granger causality. *Proc. Natl. Acad. Sci. U. S. A.* **101,** 9849–54 (2004).

7.	Dhamala, M., Rangarajan, G. & Ding, M. Analyzing information flow in brain networks with nonparametric Granger causality. *Neuroimage* **41,** 354–62 (2008).

8.	Haufe, S., Nikulin, V. V, Müller, K.-R. & Nolte, G. A critical assessment of connectivity measures for EEG data: a simulation study. *Neuroimage* **64,** 120–33 (2013).

9.	Oswal, A. *et al.* Deep brain stimulation modulates synchrony within spatially and spectrally distinct resting state networks in Parkinson’s disease. *Brain* **139,** 1482–1496 (2016).

10.	Sherman, M. A. *et al.* Neural mechanisms of transient neocortical beta rhythms: Converging evidence from humans, computational modeling, monkeys, and mice. *Proc. Natl. Acad. Sci. U. S. A.* **113,** E4885–E4894 (2016).

11.	Burke, J. F., Ramayya, A. G. & Kahana, M. J. Human intracranial high-frequency activity during memory processing: Neural oscillations or stochastic volatility? *Current Opinion in Neurobiology* **31,** 104–110 (2015).

12.	Lozano-Soldevilla, D., Huurne, N. & Oostenveld, R. Neuronal oscillations with non-sinusoidal morphology produce spurious phase-to-amplitude coupling and directionality. *Front. Comput. Neurosci.* **10,** (2016).

13.	Cole, S. R. *et al.* Nonsinusoidal beta oscillations reflect cortical pathophysiology in parkinson’s disease. *J. Neurosci.* **37,** 4830–4840 (2017).

14.	Cole, S. R. & Voytek, B. Brain Oscillations and the Importance of Waveform Shape. *Trends in Cognitive Sciences* **21,** 137–149 (2017).

15.	Vaz, A. P., Yaffe, R. B., Wittig, J. H., Inati, S. K. & Zaghloul, K. A. Dual origins of measured phase-amplitude coupling reveal distinct neural mechanisms underlying episodic memory in the human cortex. *Neuroimage* **148,** 148–159 (2017).

16.	Tinkhauser, G. *et al.* The modulatory effect of adaptive deep brain stimulation on beta bursts in Parkinson’s disease. *Brain* **140,** 1053–1067 (2017).

17.	Huang, N. E. *et al.* The empirical mode decomposition and the Hubert spectrum for nonlinear and non-stationary time series analysis. *Proc. R. Soc. A Math. Phys. Eng. Sci.* **454,** 903–995 (1998).

18.	Dayan, P. & Abbott, L. *Theoretical neuroscience: computational and mathematical modeling of neural systems*. (MIT Press, 2001).

19.	Moran, R. J. *et al.* A neural mass model of spectral responses in electrophysiology. *Neuroimage* **37,** 706–720 (2007).

20.	Nambu, A. Somatotopic Organization of the Primate Basal Ganglia. *Front. Neuroanat.* **5,** (2011).

21.	Johnson, M. D. & McIntyre, C. C. Quantifying the neural elements activated and inhibited by globus pallidus deep brain stimulation. *J. Neurophysiol.* **100,** 2549–2563 (2008).

22.	Buckwar, E. Introduction to the numerical analysis of stochastic delay differential equations. *J. Comput. Appl. Math.* **125,** 297–307 (2000).

23.	Higham, D. J. *An Algorithmic Introduction to Numerical Simulation of Stochastic Differential Equations*. *Society for Industrial and Applied Mathematics* **43,** 525–546 (2001).

24.	Pavlides, A., Hogan, S. J. & Bogacz, R. Computational Models Describing Possible Mechanisms for Generation of Excessive Beta Oscillations in Parkinson’s Disease. *PLoS Comput. Biol.* **11,** e1004609 (2015).

25.	Litvak, V. *et al.* Resting oscillatory cortico-subthalamic connectivity in patients with Parkinson’s disease. *Brain* **134,** 359–74 (2011).

26.	Fogelson, N. *et al.* Different functional loops between cerebral cortex and the subthalmic area in Parkinson’s disease. *Cereb. cortex* **16,** 64–75 (2006).

27.	Holgado, A. J. N., Terry, J. R. & Bogacz, R. Conditions for the generation of beta oscillations in the subthalamic nucleus-globus pallidus network. *J. Neurosci.* **30,** 12340–52 (2010).

28.	Van Ede, F., Quinn, A. J., Woolrich, M. W. & Nobre, A. C. Neural Oscillations: Sustained Rhythms or Transient Burst-Events? *Trends in Neurosciences* **41,** 415–417 (2018).

29.	Baker, A. P. *et al.* Fast transient networks in spontaneous human brain activity. *Elife* **2014,** (2014).

30.	Goetz, C. G. *et al.* Movement Disorder Society-sponsored revision of the Unified Parkinson’s Disease Rating Scale (MDS-UPDRS): scale presentation and clinimetric testing results. *Mov. Disord.* **23,** 2129–70 (2008).

31.	Folstein, M. F., Folstein, S. E. & McHugh, P. R. ‘Mini-mental state’. A practical method for grading the cognitive state of patients for the clinician. *J. Psychiatr. Res.* **12,** 189–198 (1975).

32.	Lambert, C. *et al.* Confirmation of functional zones within the human subthalamic nucleus: patterns of connectivity and sub-parcellation using diffusion weighted imaging. *Neuroimage* **60,** 83–94 (2012).

33.	Shink, E. & Smith, Y. Differential synaptic innervation of neurons in the internal and external segments of the globus pallidus by the GABA- and glutamate-containing terminals in the squirrel monkey. *J. Comp. Neurol.* **358,** 119–141 (1995).

34.	Nambu, A. Globus pallidus internal segment. *Progress in Brain Research* **160,** 135–150 (2007).

35.	Parent, M. & Parent, A. The pallidofugal motor fiber system in primates. *Park. Relat. Disord.* **10,** 203–211 (2004).

36.	DeLong, M. R. Primate models of movement disorders of basal ganglia origin. *Trends Neurosci.* **13,** 281–5 (1990).

37.	DeLong, M. R. & Wichmann, T. Circuits and circuit disorders of the basal ganglia. *Archives of Neurology* **64,** 20–24 (2007).

38.	Wichmann, T. & Delong, M. R. Functional and pathophysiological models of the basal ganglia. *Curr. Opin. Neurobiol.* **6,** 751–758 (1996).

39.	Kita, H., Tachibana, Y., Nambu, A. & Chiken, S. Balance of monosynaptic excitatory and disynaptic inhibitory responses of the globus pallidus induced after stimulation of the subthalamic nucleus in the monkey. *J. Neurosci.* **25,** 8611–8619 (2005).

40.	Devergnas, A. & Wichmann, T. Cortical potentials evoked by deep brain stimulation in the subthalamic area. *Front. Syst. Neurosci.* **5,** 30 (2011).
